# Mast cells interact directly with colorectal cancer cells to promote epithelial-to-mesenchymal transition

**DOI:** 10.1101/2025.03.19.644113

**Authors:** Rosie Lanzloth, Nicole L. Harris, Anthony M. Cannon, Mark H. Kaplan, Heather M. O’Hagan

## Abstract

Mast cells (MCs), a type of granulocytic immune cell, can be both pro– and anti-tumorigenic in colorectal cancer (CRC). We hypothesized that these contrasting findings may be in part due to differential interactions of MCs with CRC subtypes. BRAF mutant CRC uniquely contains intestinal secretory cell types. In this study, we demonstrated that MCs are enriched in BRAF mutant CRC, likely because they are recruited by factors released from cancer secretory cells. To investigate the functional consequences of MC-CRC cell interactions, we performed direct coculture experiments. We demonstrated that MCs promote epithelial-to-mesenchymal transition (EMT) in CRC cells in a calcium– and contact-dependent fashion. Furthermore, inhibiting LFA-1 and ICAM1 integrin binding reduced the coculture-induced EMT-related marker expression in CRC cells. The MC-CRC cell interaction facilitates the transfer of biological materials, including mRNA molecules, from MCs to CRC cells. This study is the first to report a contact-dependent, pro-tumorigenic role of MCs in CRC, as well as the transfer of molecules encoded by MCs to CRC cells. These findings enhance our comprehension of cell-cell communication between immune and cancer cells. Furthermore, this work suggests that targeting MC-CRC interactions, particularly through modulating integrin pathways, could offer new therapeutic strategies for aggressive CRC subtypes.

## Introduction

Colorectal cancer (CRC) is the third most common cancer diagnosed in the US and the second leading cause of cancer-related deaths in men and women combined (1). Of patients diagnosed with CRC as a primary tumor, 25% will later develop metastases (2). While early-stage CRC can be cured by surgery, metastatic CRC is difficult to treat due to a large burden of disseminated cancer cells throughout the body. Recent progress in CRC treatments has improved overall survival in patients with metastatic CRC (2). However, not all patients respond to targeted therapies in part due to the heterogeneity of CRC cells and the complexity of the tumor immune microenvironment.

BRAF mutant CRCs are highly aggressive, with many studies showing overall worse patient survival compared with other CRC subtypes (3,4). BRAF mutation is associated with microsatellite instability (MSI) and MSI CRCs are characterized by a significant enrichment in antitumor immune cells (5). However, approximately 50% of BRAF mutant CRCs are microsatellite-stable (MSS). In contrast to MSI CRCs, MSS CRCs typically lack antitumor immune cell enrichment and thus, respond poorly to immunotherapy, including immune checkpoint inhibitors (5). Further, research by Shields et al. has shown distinct differences in the composition of immune cell populations between BRAF wildtype and BRAF mutant tumors in mouse models (6). Therefore, it is crucial to investigate how the tumor immune microenvironment is regulated in MSS BRAF mutant and BRAF wild-type CRCs to find better immunotherapies for these difficult-to-treat cancers.

A recent study demonstrated that secretory cells, namely enteroendocrine and goblet cells, are enriched in BRAF mutant CRC (7). Secretory cells in the normal intestine are known to recruit and interact with immune cells, including mast cells (MCs) (8–12). Therefore, secretory cells present in BRAF mutant CRCs may affect how this subtype of CRC interacts with the tumor immune microenvironment.

MCs, tissue-resident granulocytic immune cells, play a significant role in tumorigenesis by promoting tumor growth, angiogenesis, and epithelial-to-mesenchymal transition (EMT) (13). EMT is a key process that enables cancer cells to migrate out of the primary tumor, which is an important early step in the metastatic process (14). However, the role of MCs in CRC is controversial. MCs can cause growth arrest and apoptosis of CRC cells (15,16). In contrast, additional studies demonstrated that MCs promoted CRC growth and invasion without establishing an exact mechanism (15). The dichotomy in the tumorigenic role of MCs in CRC might be explained by the use of distinct CRC subtypes in these studies. Therefore, we focused on BRAF mutant CRC because of the potential unique relationship between the secretory cells present in this CRC subtype and MCs.

In this study, we demonstrate that MCs are enriched in BRAF mutant human CRC and mouse colon tumors, and secretory cells in BRAF mutant CRC promote the migration of MCs in vitro. Additionally, we demonstrate that MCs promote EMT in CRC cells in an integrin-mediated, and contact– and calcium-dependent manner. We further find that AKT activation in both cell types is necessary for the promotion of EMT in CRC cells. Lastly, we demonstrate that molecules encoded by MCs are transferred into CRC cells. Altogether this work improves our understanding of how EMT can be induced in CRC cells while describing novel interactions between MC and CRC cells.

## Materials and Methods

### Cell lines

All cell lines were maintained in a humidified atmosphere with 5% CO2. HT-29 and SW403 cells were cultured in McCoys 5A media (Corning, #10-050-CV) and RPMI 1640 media (Corning, #10-040-CV), respectively, supplemented with 10% FBS (Corning, #35-015-CV). Both cell lines were purchased from the ATCC. The human MC line LAD2 (generously provided by Dr. Dean Metcalfe at NIH/NIAID, MD, USA) was cultured as described previously (17,18) in StemPro medium supplemented with 4 mM l-glutamine, 100 U/ml penicillin, and 0.1 mg/ml streptomycin and 100 ng/ml SCF (300-07, Peprotech). BMMCs were generated by differentiation of human cord blood CD34+ cells (70002.2, StemCell Technologies). For di_erentiation, CD34+ cells were supplemented with 100 µg/ml recombinant SCF, 100 µg/ml recombinant IL6 (200-06, Peprotech), and 30 µg/ml recombinant IL3 (200-03, Peprotech) for the first week only. BMMCs were used after 4 weeks of differentiation. All cells used in experiments were passaged fewer than 15 times.

### Coculture system and treatments

For the coculture system, HT-29 cells were seeded in McCoys 5A media 24 hours prior to the coculture. On the day of the coculture, HT-29 cells were washed once with PBS and LAD2 cells were added in a 1:1 ratio. The coculture was incubated in complete StemPro media (free from SCF) for indicated time points. At the end of the incubation, media containing the LAD2 cells was carefully removed and centrifuged at 500xg at 4°C for 5 minutes and pellets were saved for downstream applications. HT-29 cells were washed vigorously 3 times with PBS, scraped and centrifuged at 500xg at 4°C for 5 minutes. Pellets were saved for downstream applications. LAD2 cells were passaged one day prior to each experiment. For migration experiments, BMMCs and HT-29 cells were starved overnight in media lacking SCF or FBS, respectively. Reparixin (MedChemExpress, HY-15251), Calcium Ionophore (Sigma-Aldrich, C7522), BAPTA-AM (MedChemExpress, HY-100545), LY294002 (MedChemExpress, HY-10108), Borussertib (MedChemExpress, HY-122913), BIRT-377 (MedChemExpress, HY-110117), Lifitegrast (MedChemExpress, HY-19344) were solubilized in DMSO prior to treatment. Treatment dosages and durations are defined in the figure legends. For LY294002, BIRT-377, and Lifitegrast treatments, MCs were pretreated for 1h prior to the direct coculture. Because these drugs are reversible, treatments were kept during the duration of the coculture.

### Statistical Analyses

Statistical analyses for qPCR and functional assays were performed using Graphpad Prism 10. Specific statistical tests used are in the figure legends. qRT-PCR data from HT-29 cells in coculture with LAD2 cells was performed N=3 independent times. qRT-PCR data from SW403 in coculture with LAD2 cells and HT-29 cells in coculture with BMMCs were performed N=1 or N=2 independent times (with 3 independent biological replicates each). For western blots, representative images are presented with the number of independent biological replicates indicated in the figure legend.

### Immunohistochemistry (IHC)

Deidentified human samples were obtained after Institutional Review Board approval (5 BRAF mutant and 5 sex, age, and primary vs. metastatic matched non-BRAF–mutant CRC adenocarcinomas). Min and BLM mouse tumor tissue samples were obtained as done previously (6). All mouse experiments were covered under a protocol approved by the Indiana University Bloomington Animal Care and Use Committee in accordance with the Association for Assessment and Accreditation of Laboratory Animal Care International. IHC was performed and scored as indicated in the Supplementary Methods.

### Mouse MC fluorescence-activated cell sorting (FACS)

Following tissue processing, single-cell suspensions were stained with a fixable viability dye, surface markers, and intracellular markers. For surface staining, cells were stained in FACS buffer containing CD117 (c-Kit) (Biolegend, 105808) and CD129 (IL9R) (Millipore/Sigma-Aldrich, MABF2304F) antibodies for 30-60 minutes at 4°C. Following surface staining, cells were fixed using an IC fixation buffer (Invitrogen) for 20 minutes at 4°C in the dark. For intracellular staining, cells were washed and stained in permeabilization buffer for 5 minutes and 60 minutes respectively. Stained cells were washed 3x with FACS buffer and analyzed using an Attune flow cytometer. Data was processed and analyzed in Flowjo.

### Preparation of conditioned medium

To produce HT-29 cell conditioned medium, EV or KD cells were grown to 100% confluency in standard medium. Medium was replaced with serum-free medium for 48 hr. The conditioned medium was then collected, centrifuged at 500xg for 5 minutes at 4°C and used for migration assays.

### Migration assay

HT-29 cells were seeded on the membrane of the upper chamber of the transwell (8 µm PET membrane; Corning, 3422) in a 24-well culture plate. 24 hours later, LAD2 cells were incubated with HT-29 cells in a 1:1 ratio for 12 hours. At the end of the incubation, LAD2 cells were removed and HT-29 cells were washed 3 times with PBS. 20% FBS was then added to the lower chamber. 24 hours later transwell inserts were stained using Hema 3 Stat Pack (Thermo Fisher Scientific, #123–869). Migration inserts were randomized before manual quantification and the outer 5% of the inserts were not included during quantification to reduce edge-effect bias. All images were taken on an EVOS FL Auto microscope.

All MC migration experiments were carried out in a 24-well culture plates using transwells with 5µm PET membrane (Corning, 3421). EV or KD HT-29 cell conditioned media was used as the chemoattractant with unconditioned media serving as control. BMMCs (5 x 10^4 cells/ml) were starved overnight, resuspended in control medium, and migrated overnight. Quantification of the migration was done as described previously (CytoSelect™ 24-Well Cell Migration Assay from Cell BioLabs, Inc protocol). Briefly, transwells were removed, migrated cells were lysed, and DNA was labeled using CyQUANT™ NF Cell Proliferation Assay kit. Fluorescence was read with a fluorescence plate reader at 480 nm/520 nm. Experiments were performed in triplicates and repeated at least 3 times.

### Secreted factor analysis

Conditioned media from EV and ATOH1 KD HT-29 cells were collected (N=3 per condition), centrifuged at 500xg for 5 minutes at 4°C, and analyzed for cytokine and chemokine levels using the human cytokine/chemokine 96-Plex Discovery Assay® Array (HD96) (Eve Technologies, Calgary, AB).

### Immunofluorescence and imaging

For vimentin immunofluorescence, HT-29 cells were seeded on No 1.5 coverslip. The next day, HT-29 cells were labeled with Vybrant™ DiI Cell-Labeling Solution (Invitrogen, V22885) per manufacturer’s instructions and incubated with LAD2 cells in 1:1 ratio for 3 hours. Then, HT-29 cells were washed vigorously 3 times with PBS and stained with anti-vimentin (CST #5741, 1:50) according to manufacturer’s protocol. Images were acquired on a Leica STERLLARIS 8 Falcon scanning confocal system with MDi8-inverted microscope with LASX software (Leica Microsystems). All the images were taken at 63X magnification, 1.2A water immersion at room temperature, and processed using ImageJ.

### Live cell Imaging

HT-29 cells were seeded on Poly-D-lysine coated Nunc™ Glass Bottom Dishes (Thermo Fischer Scientific, 150680). The next day, unlabeled or DiO (10 µg/ml) (Invitrogen, D275) labeled LAD2 cells or BMMCs were incubated with HT-29 cells or SW403 cells in HEPES Tyrode’s buffer immediately upon imaging. Images were acquired on an Olympus OSR spinning disk confocal system with IX83 inverted microscope with CellSense acquisition software. All the images were taken at 40X or 60X magnification, 0.95A water immersion and 1.3A silicon immersion, respectively, at 37°C and 5% Co2, and processed using ImageJ.

### RNA isolation and RT-qPCR

Total RNA was isolated from cell pellets using the RNeasy mini kit (Qiagen #74104) according to the manufacturer’s protocol. The Maxima first strand cDNA synthesis kit (Thermo Fisher #K1642) was used to synthesize cDNA. For quantitative reverse transcription PCR, cDNA was amplified using gene-specific primers and FastStart Essential DNA Green Master (Roche #06402712001) (19). Cq values of genes of interest were normalized to housekeeping gene *RHOA* expression. Primer sequences are listed in the Supplementary Methods.

### Knockdown and plasmids

See the Supplementary Methods for details on shRNAs and plasmids used. Lentivirus was generated described previously (19).

### MC spinoculation

5×10^5 MCs were resuspended in 200 µl of fresh SP34 media and 50 µl of Vimentin-Flag viral suspension with 8 µg/ml polybrene in 12-well plates. Cells were spun at 1500xg for 90 minutes at 30°C. Fresh media was added to transduced cells. The next day, cells were resuspended in fresh media and were used for the experiments.

## Results

### MCs are enriched in BRAF mutant CRC

Based on the phenotypic and cell composition differences between BRAF-mutant and BRAF-wild-type CRC, we wanted to determine differences in immune cell composition between the two tumor types. In a previous study that compared the tumor immune microenvironment of BRAF mutant (*Braf^V600E^Lgr5^Cre^Min;* BLM) and BRAF wildtype (Min) mouse colon tumors, CIBERSORT analysis of RNA-sequencing data revealed an enrichment for MCs in BRAF mutant tumors (6). Furthermore, the expression of MC-specific proteases was significantly higher in BLM mutant than Min tumors (Figure 1A). Confirming these results, a higher number of mast cell protease 1 (MCPT1) positive cells was observed in BLM tumors compared to Min tumors by IHC (Figure 1B, 1C). FACS analysis of these tumors also indicated an enrichment of MCs in BLM tumors compared to Min tumors and the normal colon epithelium (Figure 1D). To determine if MCs were also enriched in BRAF-mutant human CRC, we performed CIBERSORT (20) analysis on The Cancer Genome Atlas (TCGA) colon adenocarcinoma (COAD) (21) patient data. There was a trend toward increased activated MCs in BRAF mutant compared to BRAF wildtype CRCs (p-value= 0.0660, Figure 1E). Similarly, there was an increased number of tryptase-positive MCs in BRAF mutant compared to BRAF wildtype human CRC samples by IHC (Figure 1F, 1G). Together this data suggests that MCs are enriched in BRAF mutant compared to BRAF wildtype CRC.

**Figure 1.**
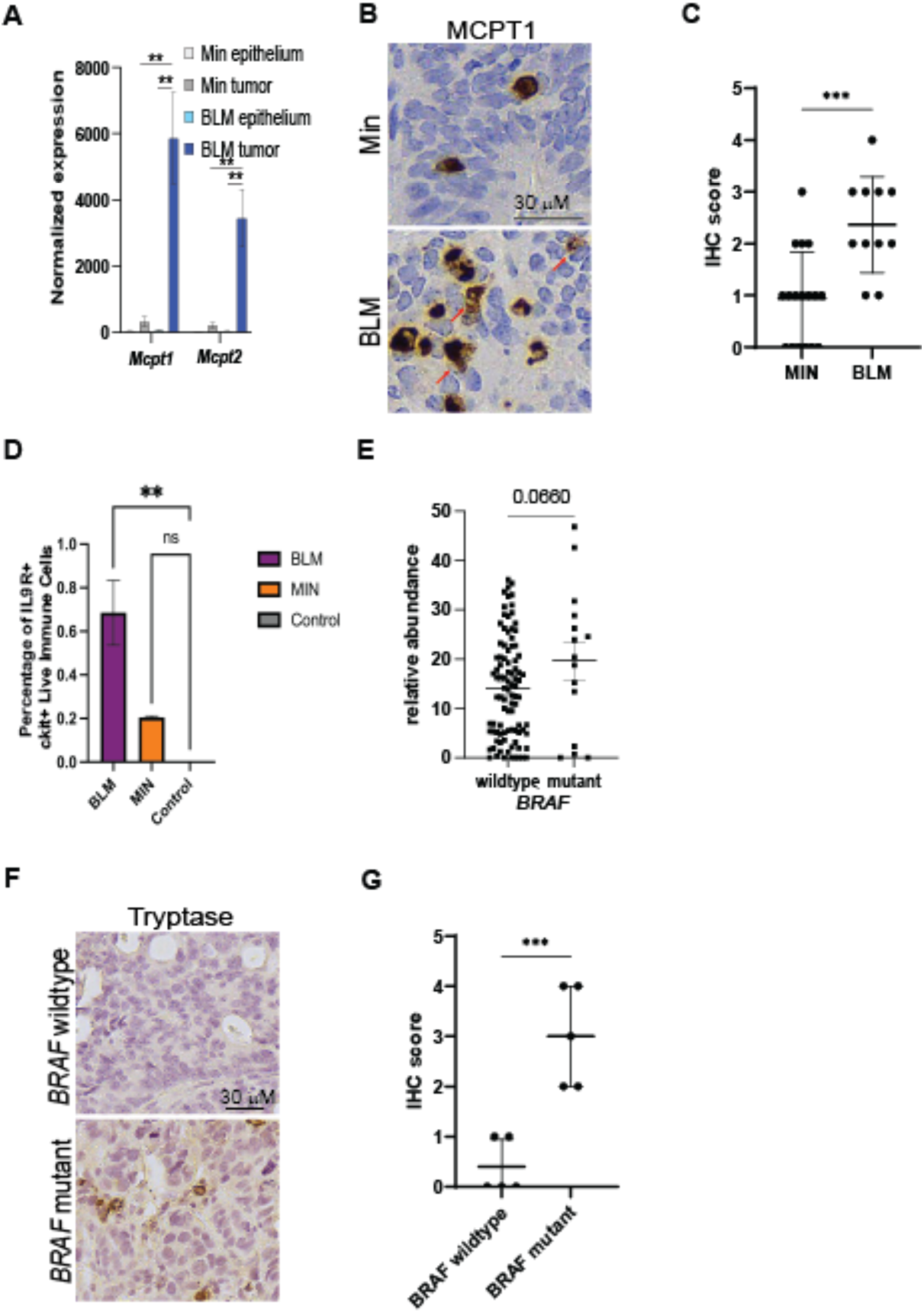
Mast cells are enriched in BRAF mutant compared to BRAF wildtype CRC. **A**. MCPT1 and MCPT2 normalized RNA-sequencing gene expression in Min and BLM colon epithelium (N=4) and colon tumors (N=4-5). **B**. IHC for MCPT1 on mouse colon Min and BLM tumors. Red arrows indicate degranulated MC. **C**. Quantification of MCPT1 positive cells from B. Each dot represents an individual tumor. Lines indicate mean −/+ SD. C. Percentage of IL9R+, ckit+ positive immune cells in BLM tumors, MIN tumors, and normal colon epithelium (control). **D**. Relative abundance of activated MCs in wildtype and BRAF mutant CRC determined by CIBERSORT analysis of RNA-sequencing data from the COAD TCGA database. Each dot represents an individual CRC. Lines indicate mean −/+ SEM. **E**. IHC for tryptase on primary human colon cancer samples. **F**. Quantification of MCPT1 positive cells from F. Graphed as in C. For all panels, significance was determined by two-tailed t-test, \**p* ≤ 0.05; **p ≤ 0.01; ***p≤ 0.001; ****p≤ 0.0001, ns-not significant.

### CRC secretory cells recruit MCs by secreting IL8

Types of secretory cells that are enriched in BRAF mutant CRC (7), including enteroendocrine cells (EECs), and goblet cells can interact with MCs (8–12) in the normal intestine, suggesting that CRC secretory cells may influence the recruitment of MCs in CRC. To address this potential connection, we differentiated CD34+ hematopoietic stem cells into bone marrow-derived mast cells (BMMCs) (Supplementary Figure S1A). Secretory cell populations were depleted from a BRAF mutant CRC cell line, HT-29, using shRNA against *ATOH1* (*7*), a transcription factor necessary for the differentiation of colon progenitor cells to secretory progenitor cells (22) (Figure 2A, Supplementary Figure S1B).

**Figure 2.**
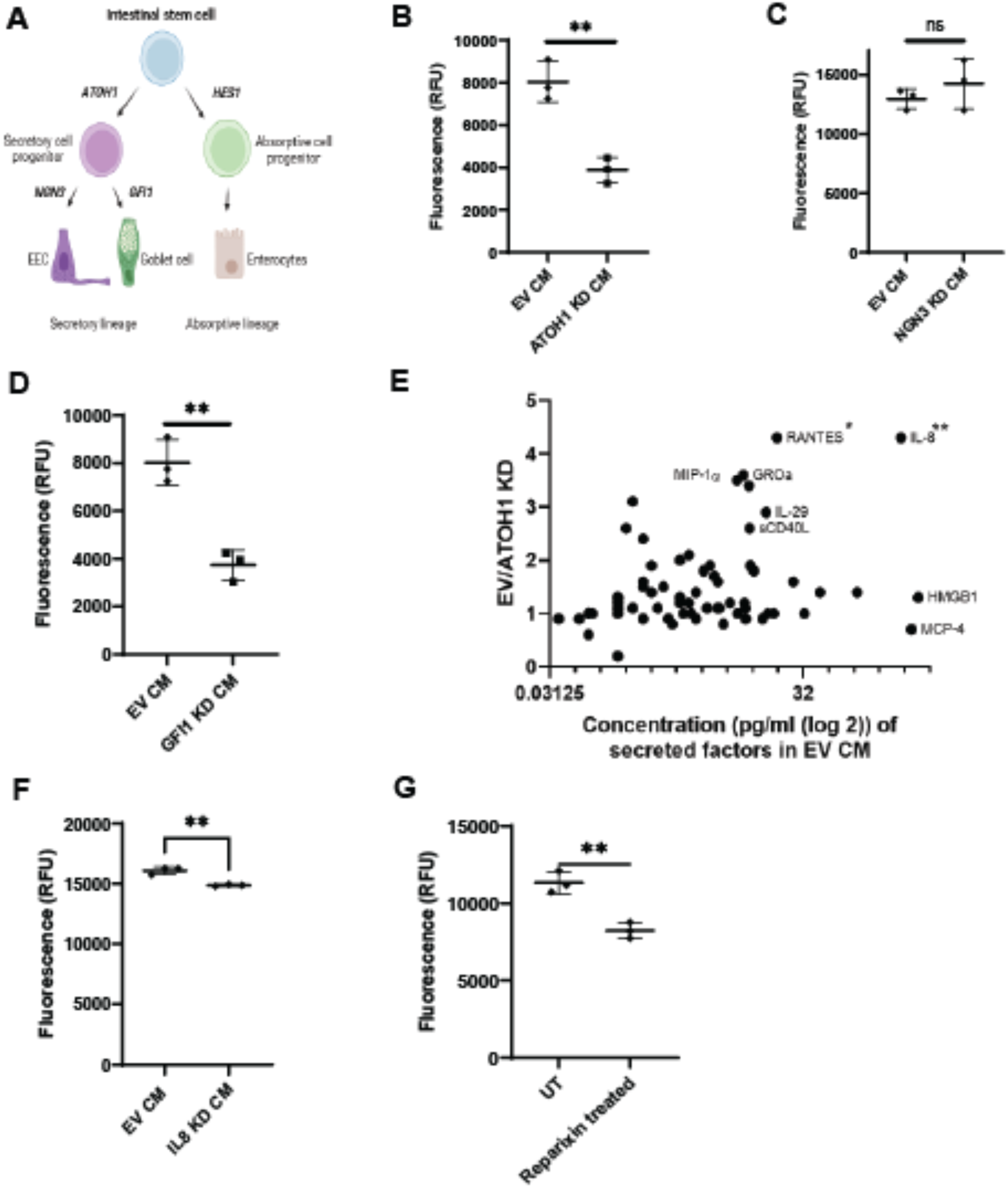
Secretory cells recruit MCs toward CRC-conditioned media by secreting IL8. **A**. Diagram of normal colon epithelial cell di_erentiation where stem cells di_erentiate down the secretory or absorptive lineages. Transcription factors required for differentiation are indicated. EEC-enteroendocrine cell. **B**. Migration of bone marrow-derived mast cells (BMMCs) toward conditioned media (CM) collected from empty vector (EV) or ATOH1 knockdown (KD) HT-29 cells assayed by quantification of DNA in the lower chambers of transwells. RFU = Relative Fluorescence Unit. Lines indicate mean +/− SD and each point represents an independent biological replicate. **C**. Migration of BMMCs toward CM of EV or GFI1 KD HT-29 cells. Data presented as in B. **D**. Migration of BMMCs toward CM of EV or NGN3 KD HT-29 cells. Data is presented as in B. **E**. Differences in factors secreted (pg/ml) by EV and ATOH1 KD HT-29 cells. EV/ATOH1 KD = values from EV CM compared to values from ATOH1 KD CM. Asterisk next to cytokine names indicates statistical significance. **F**. Migration of BMMCs toward CM of EV or IL8 KD HT-29 cells. Data is presented as in B. **G**. Migration assay of untreated (UT) or Reparixin-treated (10 µM, overnight) BMMCs toward HT-29 CM. Data is presented as in B. For all panels, N=3 and significance was determined by two-tailed t-test, \**p* ≤ 0.05; **p ≤ 0.01; ***p≤ 0.001; ****p≤ 0.0001, ns-not significant.

There was a significant reduction in the migration of BMMCs toward the conditioned media collected from ATOH1 knockdown (KD) versus empty vector (EV) HT-29 cells (Figure 2B). *NEUROG3* and *GFI1* are transcription factors required for the differentiation of EECs and goblet cells, respectively (23) (Figure 2A, Supplementary Figure S1C, S1D). Depletion of EECs by knocking down *NEUROG3* did not result in a significant difference in the migration of BMMCs (Figure 2C). However, the depletion of goblet cells by knocking down *GFI1* significantly reduced the migration of BMMCs toward conditioned media (Figure 2D).

Cytokines such as SCF are known to recruit MCs to sites of inflammation and secretory cells present in BRAF mutant CRC likely contribute to the CRC secretome (15,24). Cytokine analysis revealed that IL8 was the most concentrated cytokine in EV HT-29 cell conditioned media and the most decreased cytokine in ATOH1 KD compared to EV HT-29 cell conditioned media (Figure 2E). To evaluate the role of IL8 in the recruitment of BMMCs, we first prevented the release of IL8 into the media by knocking down *CXCL8* (IL8) in HT-29 cells (Supplementary Figure S1E). There was a significant reduction in the migration of BMMCs toward the conditioned media of IL8 KD versus EV HT-29 cells (Figure 2F). Additionally, Reparixin was used to inhibit the BMMC-expressed IL8 receptors, CXCR1, and CXCR2. There was a significant reduction in the migration of Reparixin-treated BMMCs compared to untreated BMMCs to conditioned media from HT-29 cells (Figure 2G).

Together, these results suggest that the release of IL-8 by CRC secretory cells contributes to MC recruitment in vitro.

### MCs promote epithelial-to-mesenchymal transition in CRC cells

To determine the tumorigenic role of MCs in CRC, we performed direct coculture of HT-29 cells with LAD2 cells, a human mast cell line expressing FcεRI and mast cell proteases (17) (Supplementary Figure S2A). To verify proper separation of LAD2 and HT-29 cells after direct coculture, cocultured cells were separated, labeled with CD45-AF488, an immune marker, and EpCam-AF647, an epithelial marker, and then analyzed by flow cytometry.

LAD2 and HT-29 cell fractions were more than 99% and 98% pure, respectively (Supplementary Figure S2B, S2C, S2D, S2E). Interestingly, gene and protein expression of EMT-related markers: Slug (*SNAI2),* vimentin (*VIM)*, and ZEB2 (*ZEB2)* increased in HT-29 cells after coculture with LAD2 cells (Figure 3A, 3B). The cytokine CCL2 also increased in cocultured HT-29 cells and is included as an internal control throughout the rest of the study. Similar results for *VIM, CCL2,* and *ZEB2* expression were observed in HT-29 cells cocultured with BMMCs (Supplementary Figure S2F). SW403 cells are a BRAF wildtype mucinous CRC cell line that contains secretory cells (19). Expression of the EMT-related markers also showed an upward trend in SW403 cells when co-cultured with LAD2 cells compared to the control (Supplementary Figure S2G). Increased vimentin protein expression in cocultured HT-29 cells was further confirmed by immunofluorescence (Figure 3C). EMT-related marker expression is often associated with increased migration (25). We observed a significant increase in the number of migrated HT-29 cells when they were pre-incubated with LAD2 cells compared to those that were not (Figure 3D, 3E). Together, these findings suggest that MCs promote the EMT process in CRC cells.

**Figure 3.**
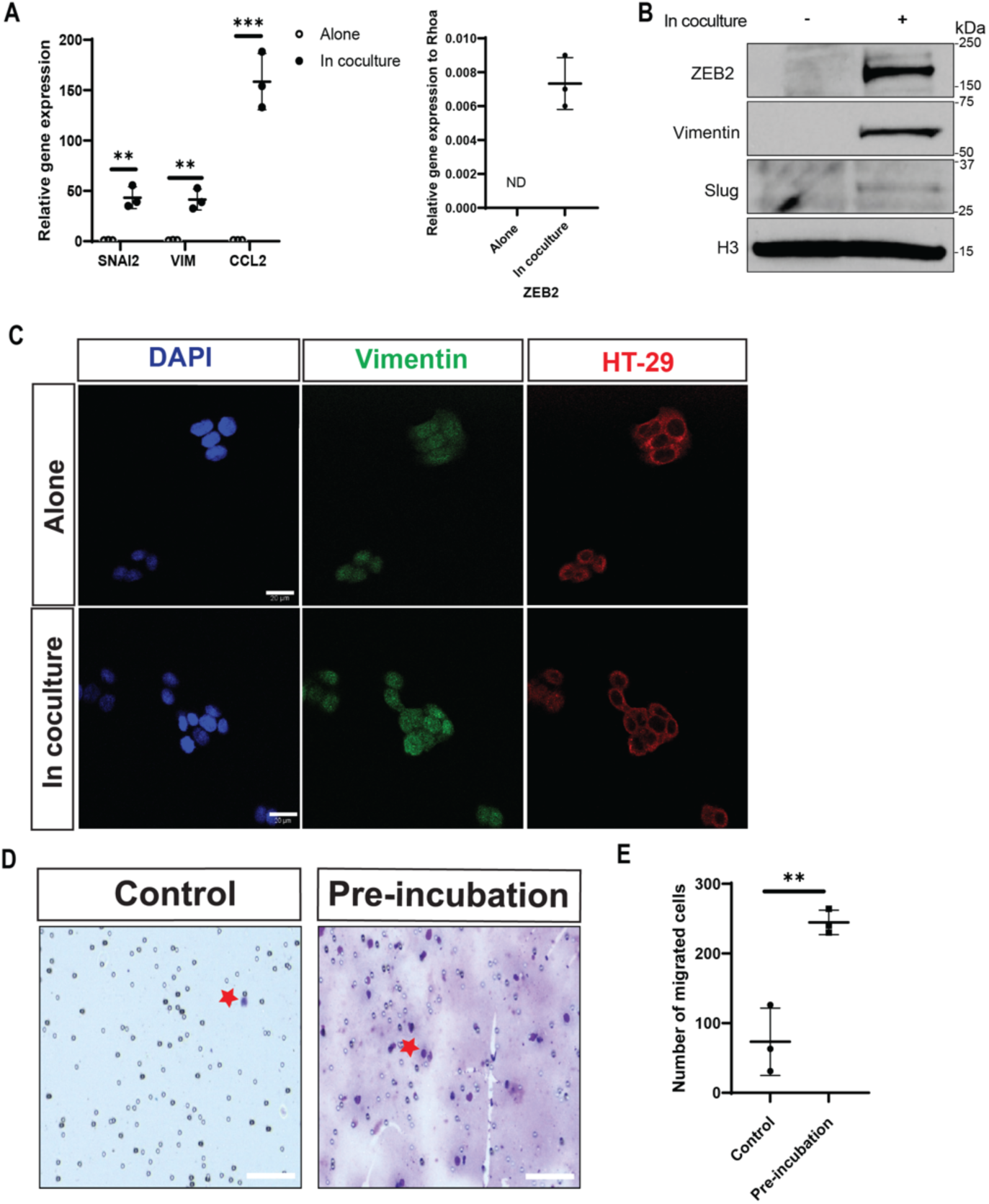
MCs induce the expression of epithelial-to-mesenchymal (EMT) related factors and promote a migratory phenotype in CRC cells. **A**. Relative gene expression of EMT markers and CCL2 (left) and ZEB2 (right, not detected (ND)) in HT-29 cells cultured alone or with LAD2 cells for 6h. **B**. Western Blot of HT-29 cells cocultured alone or with LAD2 cells for 3h. Representative blot from N=3 biological replicates. **C**. Representative immunofluorescence images of vimentin (green) in HT-29 cells (red, labeled with DiL, 10 µg/ml, 20 minutes) alone or after direct coculture with LAD2 cells for 3h. **D**. Images of membrane showing migrated HT-29 cells (indicated by red stars) that were preincubated alone (control) or with LAD2 cells (12h) after 24h. Scale bar is 100 µM. **E**. Number of migrated HT-29 cells from D. N=3. For all panels, lined represent mean +/− SD. Significance was determined by two-tailed t-test, \**p* ≤ 0.05; **p ≤ 0.01; ***p≤ 0.001; ****p≤ 0.0001, ns-not significant.

### MCs play an active role in the promotion of the EMT process in CRC cells

Calcium signaling is involved in mediator release, signal transduction, and activation in MCs (26,27). The calcium chelator, BAPTA-AM, significantly blocked mediator release in LAD2 cells (Supplementary Figure S3A). Consistent with our previous results, coculture of HT-29 and LAD2 cells increased the expression of EMT-related marker genes and proteins. However, pre-treating LAD2 cells with BAPTA-AM blocked the induction of EMT-related markers in cocultured HT-29 cells (Figure 4A, 4B). Similar trends were observed when BAPTA-AM pretreated BMMCs were cocultured with HT-29 cells, and when SW403 cells were cocultured with BAPTA-AM pretreated LAD2 cells (Supplementary Figure S3B, S3C, S3D, S3E). BAPTA-AM pre-treatment of LAD2 cells did not affect Vimentin levels in LAD2 cells, further confirming the absence of contamination from the LAD2 cell fraction to the cancer cell fraction (Supplementary Figure S3F). No significant differences were observed in the expression of the EMT-related genes in BAPTA-AM pretreated HT-29 cells compared to non-pretreated cocultured HT-29 cells (Supplementary Figure S3G) suggesting that calcium signaling in HT-29 cells was not involved in the induction of EMT-related markers.

**Figure 4.**
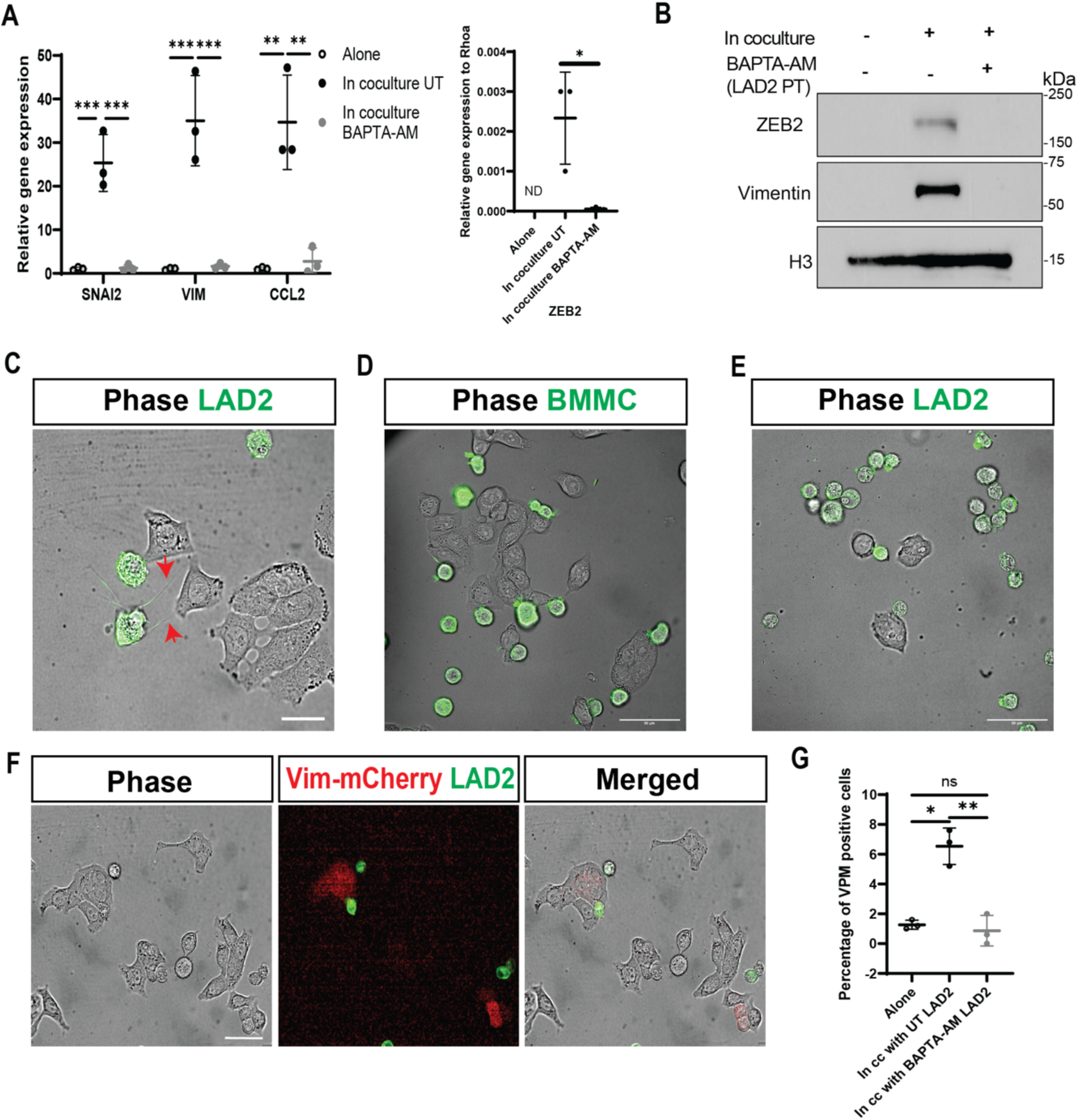
MCs induce EMT in CRC cells in a calcium– and contact-dependent manner. **A**. Relative qRT-PCR of EMT genes and CCL2 (left) and ZEB2 (right, not detected (ND)) in HT-29 cells alone, in coculture with unpretreated (UT) LAD2 cells, or in coculture with BAPTA-AM (20 µM, 1h) pretreated LAD2 cells for 6h. **B**. Western blot of HT-29 cells alone, in coculture with unpretreated LAD2 cells, or in coculture with BAPTA-AM (20 µM, 1h) pretreated LAD2 cells (LAD2 PT) for 3h. N=3. **C**. LAD2 cells were labeled with DiO (10 µg/ml, 20 minutes, green) and incubated in direct coculture with HT-29 cells (unlabeled). Live cells were imaged with Olympus OSR SD confocal microscope for 4h. Arrowheads indicate cytoplasmic extensions. Magnification 100x. Scale bar is 20 µM. **D**. BMMCs were incubated in direct coculture with HT-29 cells. Cells were labeled and imaged for 1h as in C. Magnification 60x. **E**. LAD2 cells were incubated in direct coculture with SW403 cells. Cells were labeled and imaged for 1h as in C. Magnification 60x. **F**. HT-29 cells transfected with a plasmid containing the vimentin promoter driving expression of mCherry were incubated with DiO labeled (10 µg/ml, 20 minutes, green) LAD2 cells. Cells were imaged as in C for 4h. Magnification 40x. Scale bar is 50 µM. **G**. Percentage of mCherry positive HT-29 cells alone, in coculture (in cc) with unpretreated (UT) LAD2 cells or in coculture with BAPTA-AM (20 µM, 1h) pretreated LAD2 cells. VPM= Vimentin promoter driving mCherry gene. For all panels, lines indicate mean +/− SD and each point represents an independent biological replicate. Significance was determined by two-tailed t-test (A, right) and one-way ANOVA (A left, and G), \**p* ≤ 0.05; **p ≤ 0.01; ***p≤ 0.001; ****p≤ 0.0001, ns-not significant.

MCs carry out their functions mainly via the release of soluble mediators (28–32). The incubation of HT-29 cells in normal or concentrated cocultured conditioned media did not result in significant differences in the expression of EMT-related genes compared to HT-29 cells incubated alone (Supplementary Figure S3H, S3I) suggesting MC-secreted factors were not sufficient to promote EMT.

Interestingly, live cell imaging of LAD2 and HT-29 cell coculture revealed direct interactions between the cell types, including direct cell-to-cell contact and cytoplasmic extensions emerging from LAD2 cells (Figure 4C). Similar observations were made between BMMCs and HT-29 cells as well as between LAD2 and SW403 cells (Figure 4D, 4E, Supplementary Figure S3I, Supplementary Video SV1). HT-29 cells transfected with a plasmid containing a Vimentin-promoter driving the expression of the mCherry gene were cocultured with LAD2 cells labeled in green. mCherry-positive clusters of HT-29 cells were found in direct contact with and in close vicinity of LAD2 cells (Figure 4F). Approximately 7% of HT-29 cells were positive for the expression of mCherry when they were cocultured with LAD2 cells compared to approximately 1% of HT-29 cells incubated alone (Figure 4G). When HT-29 cells were cocultured with BAPTA-AM pretreated LAD2 cells the number of mCherry-positive HT-29 cells was similar to HT-29 cells incubated alone (Figure 4G). Together, these findings suggest that MCs play an active role in the induction of EMT-related marker expression in CRC cells in a contact-dependent manner.

### PI3K/AKT pathway inhibition leads to decreased EMT-related expression in cocultured CRC cells

BAPTA-AM pre-treatment of LAD2 cells affected their morphology but did not prevent the direct interactions with HT-29 cells from occurring (Supplementary Figure S4A, S4B, S4C, S4D). Interestingly, AKT activation was decreased in LAD2 cells pretreated with BAPTA-AM compared to the control (Supplementary Figure S4E) leading us to hypothesize that AKT activity is important for the MC-induced EMT in CRC cells.

LY294002 (reversible PI3K inhibitor) treatment of the direct coculture reduced the gene and protein expression of most of the EMT-related markers related to untreated cocultured HT-29 cells (Figure 5A, 5B). As expected, AKT activation was reduced in cocultured HT-29 cells under LY294002 treatment. LY294002 treatment also reduced the coculture-induced increase in EMT-related marker expression in HT-29 cells in coculture with BMMCs (Figure 5C, 5D). Similarly, EMT-related markers were decreased in SW403 cells cocultured with LAD2 cells under LY294002 treatment (Supplementary Figure S4F, S4G).

**Figure 5.**
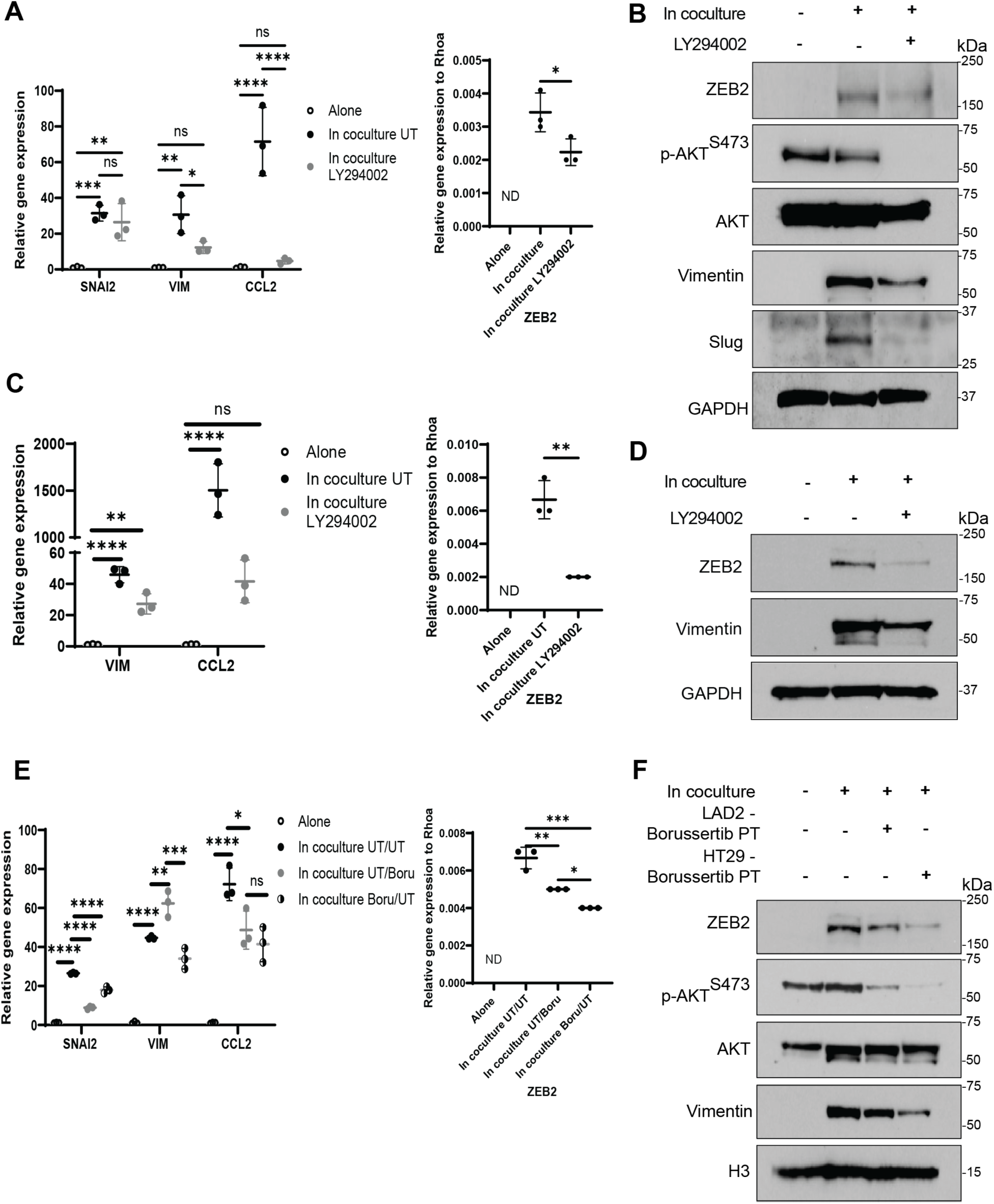
AKT activation is involved in the cocultured-induced increase of EMT-related marker expression in CRC cells. **A**. Relative qRT-PCR of EMT related genes and CCL2 (left) and ZEB2 (right, not detected (ND)) in HT-29 cells alone, in coculture with untreated (UT) LAD2 cells, or in coculture with LAD2 cells under LY294002 treatment (50 µM). **B**. Western blot of HT-29 cells treated and cocultured as in A. N=3. **C**. Relative qRT-PCR of EMT related genes and CCL2 (left) and ZEB2 (right, not detected (ND)) in HT-29 cells alone, in coculture with untreated (UT) BMMCs, or in coculture with BMMCs under LY294002 treatment (60 µM). **D**. Western blot of HT-29 cells treated and cocultured as in C for 3h. N=1. **E**. Relative qRT-PCR of EMT related genes and CCL2 (left) and ZEB2 (right, not detected (ND)) in HT-29 cells alone, unpretreated and in coculture with unpretreated LAD2 cells (UT/UT), unpretreated and in coculture with Borussertib (1 µM, overnight) pretreated (PT) LAD2 cells (UT/Boru) or Borussertib (10 µM, overnight) pretreated in coculture with unpretreated LAD2 cells (Boru/UT) for 6h. **F**. Western blot of HT-29 cells treated and cocultured as in E for 3h. HT-29 – Borussertib PT = Borussertib pretreated HT-29 cells. LAD2 – Borussertib PT = Borussertib pretreated LAD2 cells. N=3. For all panels, lines indicate mean +/− SD and each point represents an independent biological replicate. Significance was determined by two-tailed t-test (A right, C right) and one-way ANOVA (A left, C left, E), \**p* ≤ 0.05; **p ≤ 0.01; ***p≤ 0.001; ****p≤ 0.0001, ns-not significant.

To determine in which cell type PI3K/AKT pathway activation was required for induction of EMT-related markers, LAD2 and HT-29 cells were individually pretreated with the irreversible AKT inhibitor Borussertib. Individual Borussertib pre-treatment of LAD2 or HT-29 cells decreased the coculture-induced increase in gene and protein expression of the EMT-related markers in the cocultured HT-29 cells, except for *VIM* gene expression when HT-29 cells were pretreated with Borussertib (Figure 5E, 5F). As expected, AKT activation was decreased in HT-29 cells pretreated with Borussertib. However, Borussertib pre-treatment of LAD2 cells also reduced AKT activation in untreated HT-29 cells (Figure 5F). Borussertib treatment of HT-29 cells also reduced the levels of activated AKT in untreated LAD2 cells (Supplementary Figure S4H). Individual Borussertib pre-treatment of LAD2 and SW403 cells also reduced the coculture-induced increase in EMT-related marker gene expression in cocultured SW403 cells, except for *ZEB2* gene expression (Supplementary Figure S4I). Protein expression of ZEB2 and Vimentin in SW403 cells was still induced by the coculture when SW403 cells were pretreated with Borussertib but was blocked when LAD2 cells were pretreated (Supplementary Figure S4J). In contrast to the results with LAD2 cells and LY294002 treatment of BMMC cocultures, individual Borussertib pre-treatment of HT-29 cells and BMMCs did not reduce the coculture-induced increase in EMT-related marker expression in HT-29 cells even though AKT activation in HT-29 cells was reduced (Supplementary Figure S4K, S4L). Of note, AKT activation was barely detectable in BMMCs (Supplementary Figure S4M).

### LFA-1/ICAM-1 integrins are involved in the induction of the EMT-related marker expression in cocultured CRC cells

Lymphocyte function-associated antigen 1(LFA-1) and Intercellular adhesion molecule-1 (ICAM-1) integrin engagement has been shown to be important in the formation of natural killer cell immunological synapses with cancer cells (33). Interestingly, in gene ontology enrichment of RNA-sequencing data, the “integrin binding” pathway was significantly upregulated in LAD2 cells in coculture with HT-29 cells compared to when LAD2 cells were incubated alone (Supplementary Figure S5A). Furthermore, *ICAM-1* expression was upregulated in HT-29 cells in coculture with LAD2 cells (Supplementary Figure S5B).

Treatment with the LFA-1 inhibitor BIRT-377 decreased the gene and protein expression of EMT-related markers in HT-29 cells cocultured with LAD2 cells compared to HT-29 cells cocultured without treatment (Figure 6A, 6B). Similar results were observed in HT-29 cells cocultured with BMMCs (Figure 6C, 6D). The coculture-mediated induction of EMT-related marker expression was also decreased by BIRT-377 in SW403 cells in coculture with LAD2 cells (Supplementary Figure S5C, S5D). Interestingly, BIRT-377 treatment also decreased HT-29 cell migration compared to when HT-29 cells were pre-incubated with untreated LAD2 cells (Figure 6E).

**Figure 6.**
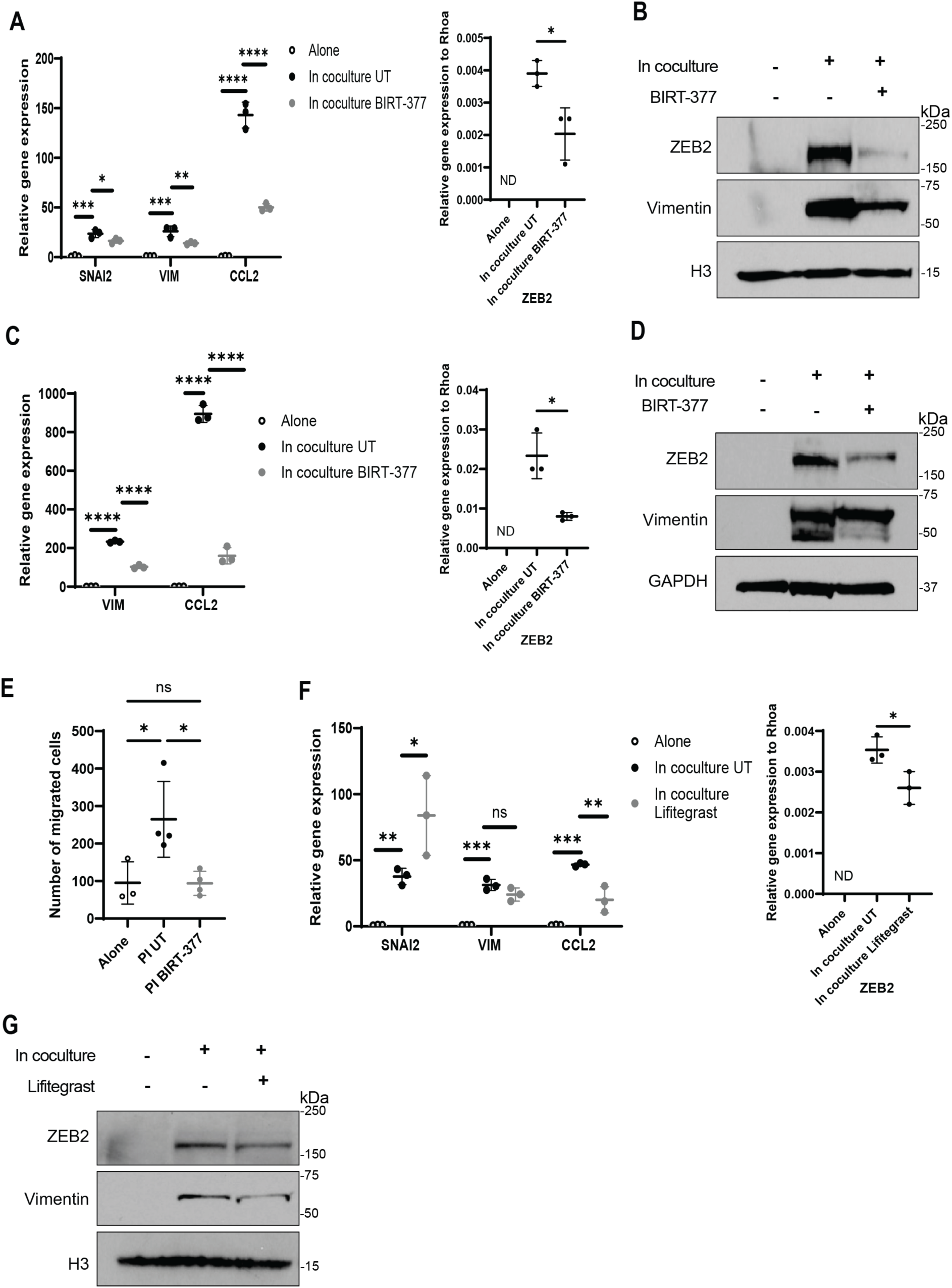
LFA-1/ICAM-1 integrin inhibition leads to a decrease in EMT-related marker expression in cocultured CRC cells. **A**. Relative qRT-PCR of EMT-related genes and CCL2 (left) and ZEB2 (right, not detected (ND)) in HT-29 cells alone, in coculture with untreated (UT) LAD2 cells, or in coculture with LAD2 cells under BIRT-377 treatment (20 µM) for 6h. **B**. Western blot of HT-29 cells treated and cocultured as in A for 3h. N=2. **C**. Relative qRT-PCR of EMT related genes and CCL2 (left) and ZEB2 (right, not detected (ND)) in HT-29 cells alone, in coculture with untreated (UT) BMMCs, or in coculture with BMMCs under BIRT-377 treatment (40 µM) for 6h. **D**. Western blot of HT-29 cells treated and cocultured as in C for 3h. N=2. **E**. Quantitative results of the migration assay of HT-29 cells preincubated (PI) alone, with untreated (PI UT) LAD2 cells, or with LAD2 cells under BIRT-377 treatment (PI BIRT-377) (20 µM) for 12h. Cells migrated for 24h. N=3. **F**. Relative qRT-PCR of EMT-related genes and CCL2 (left) and ZEB2 (right, not detected (ND)) in HT-29 cells alone, in coculture with untreated (UT) LAD2 cells, or in coculture with LAD2 cells under Lifitegrast treatment (40 µM) for 6h. **G**. Western blot of HT-29 cells treated and cocultured as in F for 3h. N=2. For all panels, lines indicate mean +/− SD and each point represents an independent biological replicate. Significance was determined by two-tailed t-test (F, right) and one-way ANOVA (A, C, E, F left) \**p* ≤ 0.05; **p ≤ 0.01; ***p≤ 0.001; ****p≤ 0.0001, ns-not significant.

Similarly to BIRT-377, treatment of the direct coculture between LAD2 and HT-29 cells with Lifitegrast, another LFA-1 inhibitor, decreased the coculture-induced increase of EMT-related markers in HT-29 cells, except for the gene expression of *SNAI2* (Figure 6F, 6G). Similar results were observed in HT-29 cells in coculture with BMMCs (Supplementary Figure S5E, S5F).

### Vimentin encoded by MCs is transferred to CRC cells

Transfer of biological materials such as organelles, vesicles, proteins, and mRNA between cells in a contact-mediated fashion is a well-established phenomenon (34–37). To determine whether LAD2 cells could transfer Vimentin molecules directly into HT-29 cells, LAD2 cells transduced with a plasmid containing Vimentin fused to FLAG (Vim-Flag) were used in direct coculture with HT-29 cells. Consistent with previous results, ZEB and Vimentin protein levels increased in HT29 cells cocultured with EV and Vim-Flag expressing LAD2 cells. Interestingly, Vim-Flag protein was detected in HT-29 cells cocultured with Vim-Flag LAD2 cells, but not EV-LAD2 cells, suggesting that material was being directly transferred from MCs to CRC cells (Figure 7A). Consistent with *VIM* and *ZEB2* protein expression, when HT-29 cells were cocultured with BAPTA-AM pretreated Vim-Flag LAD2 cells, the presence of Vim-Flag was not detected in HT-29 cells, even though Vim-Flag was still expressed in the BAPTA-AM treated Vim-Flag LAD2 cells (Figure 7A, Supplementary Figure S6A). Vim-Flag was also detected in SW403 cells in coculture with Vim-Flag LAD2 cells and was blocked by BAPTA-AM pretreatment of the LAD2 cells (Figure 7B). Similar to previous results, the LFA-1 integrin inhibitor BIRT-377 decreased the coculture-induced increase in EMT-related markers in HT-29 cells (Figure 7C). Likewise, the transfer of Vim-Flag from LAD2 cells to HT-29 cells was decreased when the direct coculture was treated with BIRT-377 compared to when the direct coculture was untreated, even though the expression of Vim-Flag did not change in BIRT-377 Vim-Flag treated LAD2 cells (Figure 7C. Supplementary Figure S6B). The transfer of Vim-Flag was also decreased in SW403 cells cocultured with Vim-Flag LAD2 cells under BIRT-377 treatment (Supplementary Figure S6C). The transfer of Vim-Flag to HT-29 cells was also decreased when the direct coculture was treated with LY294002, even though the expression of Vim-Flag did not change in LY294002 treated Vim-Flag LAD2 cells (Figure 7D, Supplementary Figure S6D). Finally, Vim-Flag mRNA was detected in HT-29 cells only when they were in coculture with Vim-Flag LAD2 cells (Figure 7E). This data suggests that LAD2 cells are transferring Vim-Flag mRNA, and possibly protein, into HT-29 cells in an integrin-mediated and calcium-dependent fashion.

**Figure 7.**
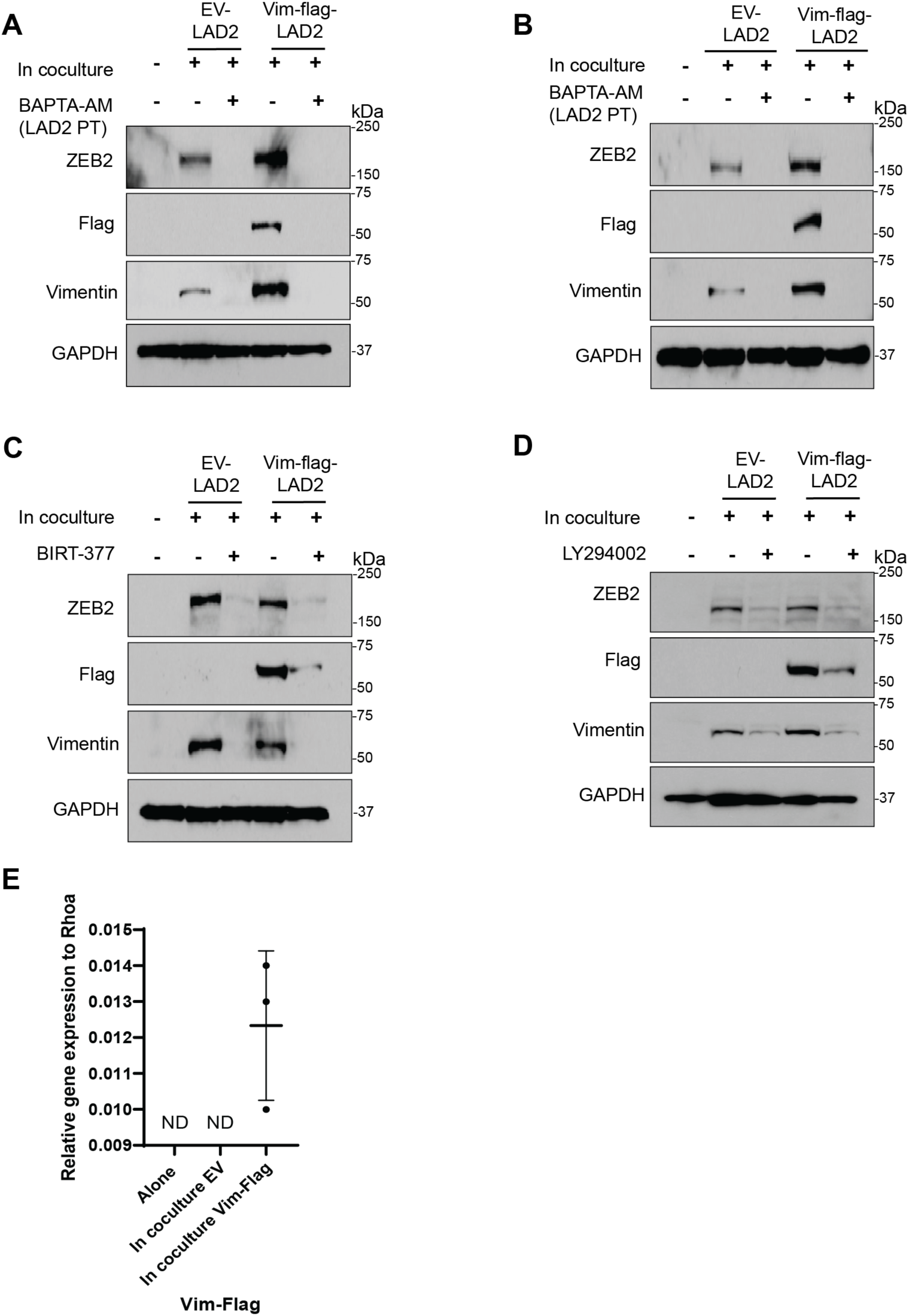
MC-encoded Vimentin-flag is transferred to CRC cells. **A**. Western blot of HT-29 cells alone or in coculture with unpretreated empty vector (EV-) LAD2 cells, BAPTA-AM (20 µM, 1h) pretreated EV LAD2 cells, unpretreated Vimentin-Flag transduced (Vim-flag-) LAD2 cells, or BAPTA-AM (20 µM, 1h) pretreated Vim-Flag LAD2 cells. N=1. **B**. Western blot of SW403 cells cocultured as in A. N=1. **C**. Western blot of HT-29 cells alone, in coculture with untreated EV LAD2 cells or in coculture with EV LAD2 cells under BIRT-377 treatment (20 µM), untreated Vim-Flag LAD2 cells, or Vim-Flag LAD2 cells under BIRT-377 treatment (20 µM). N=1. **D**. Western blot of HT-29 cells alone or in coculture with untreated EV LAD2 cells, EV LAD2 cells under LY294002 treatment (50 µM), untreated Vim-Flag LAD2 cells, or Vim-flag LAD2 cells under LY294002 treatment (50 µM). N=1. **E**. Relative qRT-PCR of Vim-Flag in HT-29 cells alone or in coculture with EV LAD2 cells, Vim-Flag transduced LAD2 cells. ND – not detected. Lines indicate mean +/− SD and each point represents an independent biological replicate.

## Discussion

In this study, we demonstrated that MCs are enriched in BRAF mutant CRC, likely because they are recruited by factors released by secretory cells. Once recruited to the tumor site, our data suggests that MCs promote the EMT process in CRC cells in an integrin-mediated and contact– and calcium-dependent fashion. Through this interaction, MCs transfer biological materials into the cancer cells, as demonstrated by the transfer of Vimentin-flag mRNA from the MCs to the CRC cells. Consistent with the EMT-related marker expression, the transfer of Vimentin-Flag mRNA is integrin-mediated and calcium-dependent. This work that demonstrates the tumorigenic role of MCs in BRAF mutant CRC has the potential to identify new therapeutic targets for these difficult-to-treat cancers.

The role of MCs in cancer remains controversial, as it varies depending on the localization of MCs within the tumor (peritumoral or intratumoral) (38) and how the MCs are activated by the tumor microenvironment (29). Therefore, the role of MCs in prognosis may differ between subtypes of CRC which have different cancer cell populations and microenvironments. Secretory cells in the normal colon are known to interact with MCs (8–12). Therefore, we hypothesized secretory cells enriched in BRAF mutant CRC (7) recruit MCs. Depleting secretory cell populations from HT-29 cells led to a reduction in BMMC migration in vitro. This result suggests that CRC secretory cells recruit MCs at least in vitro. Therefore, we hypothesize that the presence of intratumoral MCs is more likely in secretory cell-enriched CRC tumors.

MCs can promote tumor growth, angiogenesis, and EMT (13,29,31,32) but published results on the role of MCs in CRC are conflicting. For example, conditioned media from MCs have been demonstrated to cause growth arrest and apoptosis of CRC cells (16) but crosstalk between MCs and CRC cells has also been shown to promote CRC cell growth (15). Most MC functions are carried out by the release of biological factors (29,38). By performing conditioned media exchange experiments, we found that factors secreted into the media by MCs, including exosomes, were not sufficient to drive the EMT-related marker expression in CRC cells. Instead, we determined that MCs physically interact with CRC cells including by direct interaction and cytoplasmic extensions. This direct interaction is required for the MCs to induce EMT related changes in the CRC cells. The need for direct interactions for at least some of the effects of MCs on CRC cells may help explain conflicting results in the literature. Our study is the first to report a pro-tumorigenic role of MCs in a contact-dependent fashion in CRC. Direct interactions between intratumoral MCs and CRC cells may contribute to tumor growth and metastasis development.

RNA sequencing data revealed enrichment in the “integrin binding” pathway in MCs cocultured with CRC cells. Our findings suggest that MCs and CRC cells most likely bind together via LFA-1/ICAM-1 integrins in a similar manner by which MCs have been shown to bind to other immune cells (39,40). Even though LFA-1 integrin inhibitors led to a reduction in the EMT-related marker expression, it was not completely blocked. This finding implies other cell adhesion molecules may be involved in the binding of MCs to CRC cells. A more thorough understanding of the role of integrin binding of MCs to CRC cells in tumorigenesis might open a new therapeutic avenue in the treatment of CRC (41).

Using a calcium chelator, we demonstrated that MCs play an active role in the promotion of the EMT process in CRC. However, MCs were still capable of physically interacting with CRC cells under the calcium chelator pre-treatment suggesting that interaction alone is not sufficient to promote EMT in the CRC cells. Calcium signaling is involved in cytokine production, exocytosis, and transmission of signaling pathways in MCs (26,27). The chelation of calcium in MCs reduced the level of AKT activation. Further investigation is needed to clarify whether the calcium chelation-induced reduction of AKT activation in MCs affects other processes, such as exocytosis, that could contribute to the expression of EMT-related markers in CRC cells.

Many different types of biological materials can be transferred between cells in a contact-mediated fashion (38–41). Interestingly, we detected MC-encoded Vimentin-Flag mRNA in cocultured CRC cells suggesting that MCs can directly transfer mRNA to CRC cells. Whether the Vim-Flag protein that was also detected in CRC cells originates directly from MCs and/or is produced from the transferred mRNA by the translation machinery of the CRC cells is unclear. In addition to MC-encoded molecules being transferred to CRC cells, MC coculture also induced the expression of the HT29 cell-encoded EMT-related markers as demonstrated by coculture inducing expression of the HT-29 cell-encoded Vimentin-promoter mCherry reporter. Therefore, we speculate that the direct transfer of vimentin mRNA from MCs leads to a positive feedback in vimentin expression in CRC cells. Likewise, the direct transfer of EMT-related transcription factor ZEB2 might lead to the endogenous expression of vimentin in CRC cells. However, the detailed mechanism by which molecules are transferred between MCs and CRC cells and how these molecules further induce the expression of EMT markers in CRC cells requires further study.

Whether MCs promote EMT-related marker expression in tumor epithelial cells in a direct contact fashion in vivo remains unknown. Moreover, this study only focused on MC-CRC cell interactions but changes to both cell types might alter other cells present in the tumor microenvironment. For example, the cytokine CCL2 is known to recruit and activate myeloid-derived suppressor cells in CRC and other cancers (42,43). *CCL2* expression was increased in CRC cells in coculture with MCs. This finding possibly suggests that direct contact between MC and CRC cells alters the cytokine composition of the tumor microenvironment, which could result in a change in tumor immune cell populations. Further studies are warranted to investigate MC-CRC interactions in vivo and the potential effects of such interactions in the alteration of the tumor immune microenvironment.

## Acknowledgements

We would like to thank the IU Bloomington Light Microscopy Imaging Center for maintaining and providing access to imaging equipment, the IU-Bloomington Flow Cytometry Core Facility for proving instrumentation and technical advice, and the IU Center for Genomics and Bioinformatics for their assistance with library preparation and sequencing of the RNA-seq samples. We thank Dr. Dean Metcalfe for generously providing the LAD2 cells. This work was in part supported by a NIH National Cancer Institute Grant (R01CA286090) [to H.M. O’Hagan], a Research Enhancement Grant [to H. M. O’Hagan] from the Indiana University School of Medicine (IUSM), a Core Pilot Grant [to H. M. O’Hagan] from the Indiana Clinical and Translational Sciences Institute funded, in part by Grant Number UL1TR002529 from the NIH, National Center for Advancing Translational Sciences, Clinical and Translational Sciences Award, and a pilot award from the IU Simon Comprehensive Cancer Center [to M.H.K.]. Core facility usage was also supported by IU Simon Cancer Center Support Grant P30 CA082709 and U54 DK106846. The content is solely the responsibility of the authors and does not necessarily represent the official views of the IUSM.

## Author Contributions

RL was responsible for designing the study, conducting the search, extracting and analyzing data, interpreting results, writing the study, and editing. NLC was responsible for analyzing data. AC was responsible for extracting and analyzing data. MHK was responsible for conceptualization and editing the manuscript. HOH contributed to the design of the study, provided project oversight, obtained funding, and edited the manuscript

## Data Availability Statement

The datasets generated during and/or analyzed during the current study are available from the corresponding author on reasonable request.

