## Supplemental Figures S1, S2, S3, S4, S5 and S6 for "Mast cells interact directly with colorectal cancer cells to promote epithelial-to-mesenchymal transition"

### Supplementary Figures

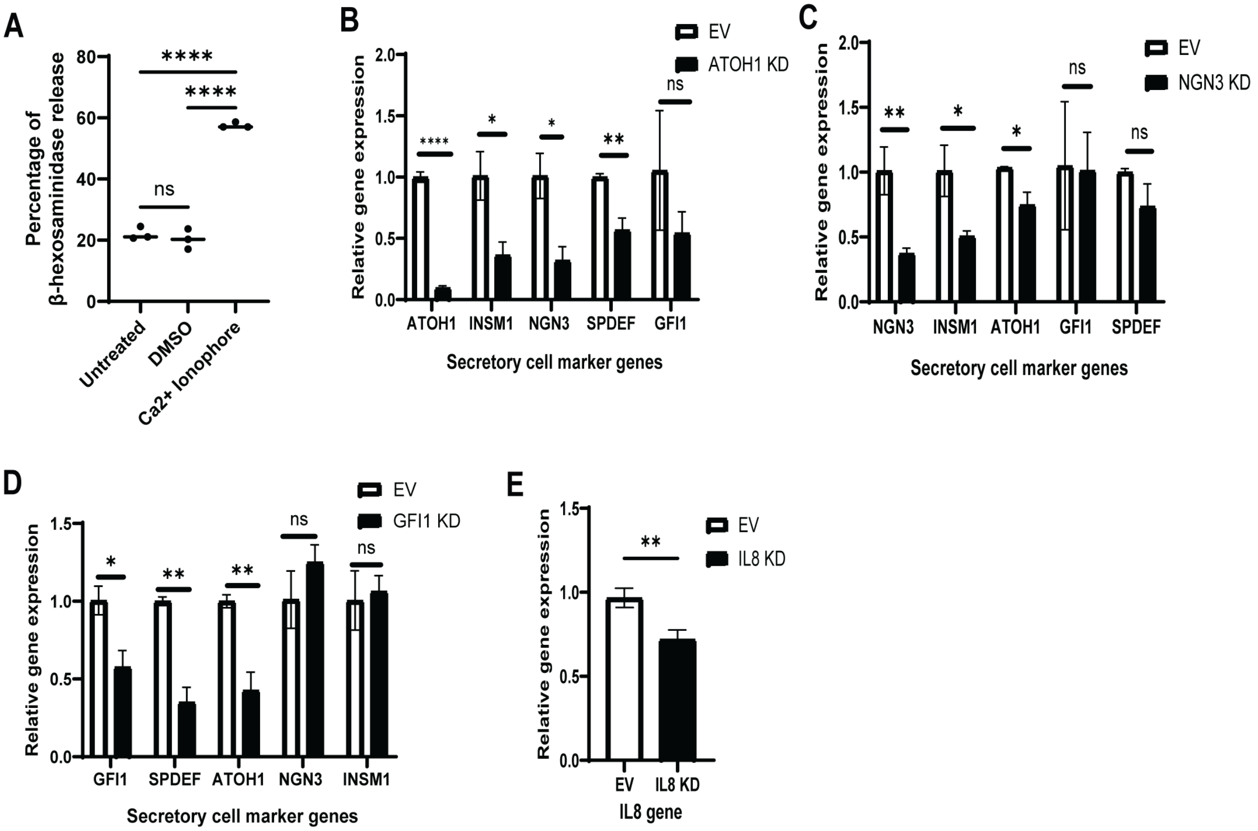


**Supplementary Figure S1. BMMC differentiation and knockdown confirmation. A**. Percentage of B-hexosaminidase released from untreated, DMSO-treated, and Calcium ionophore (Ca2+ Ionophore, 2 μM, 1h) treated BMMCs. Line indicates the mean and each point represents an independent biological replicate. **B**. Relative qRT-PCR of secretory cell marker genes in empty vector (EV) and ATOH1 knockdown (KD) HT-29 cells. **C**. Relative qRT-PCR of secretory cell marker genes in EV and GFI1 KD HT-29 cells. Data presented as in B. **D**. Relative qRT-PCR of secretory cell marker genes in EV and NGN3 KD HT-29 cells. Data presented as in B. **E**. Relative qRT-PCR of IL8 in EV and IL8 KD HT-29 cells. Data presented as in B. For all panels, graphs indicate mean +/- SD. Significance was determined by two-tailed t-test (E) and one-way ANOVA (A, B, C, and D), *$p \leq 0.05$; **$p \leq 0.01$; ***$p \leq 0.001$; ****$p \leq 0.0001$, ns- not significant.

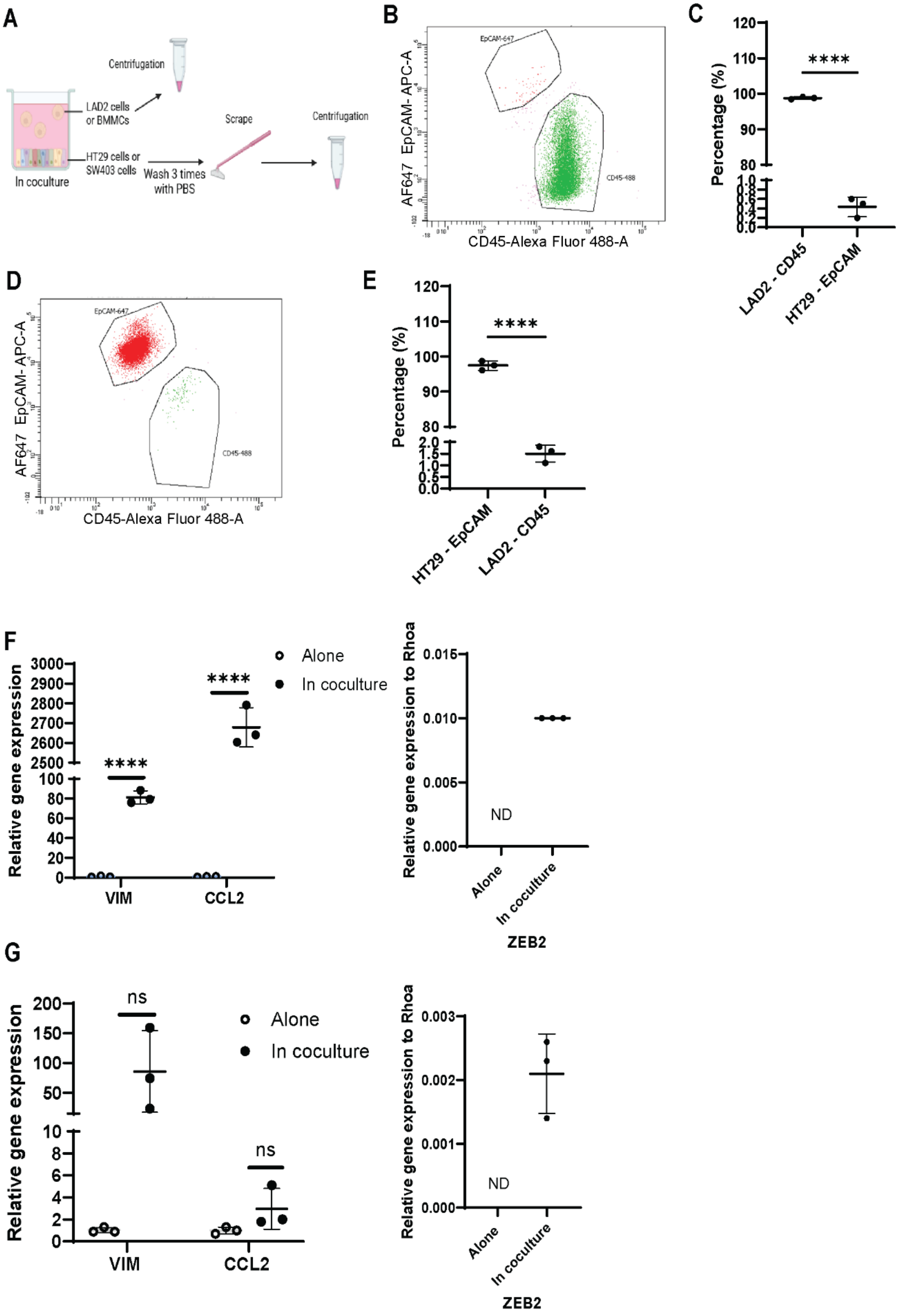
A
Centrifugation
LAD2 cells or BMMCs
HT29 cells or SW403 cells
In coculture
Wash 3 times with PBS
Scrape
Centrifugation
B
EpCAM-647
CD45-488
AF647 EpCAM- APC-A
CD45-Alexa Fluor 488-A
C
****
Percentage (%)
LAD2 - CD45
HT29 - EpCAM
D
EpCAM-647
CD45-488
AF647 EpCAM- APC-A
CD45-Alexa Fluor 488-A
E
****
Percentage (%)
HT29 - EpCAM
LAD2 - CD45
F
Relative gene expression
****
****
Alone
In coculture
VIM
CCL2
Relative gene expression to Rhoa
ND
Alone
In coculture
ZEB2
G
Relative gene expression
ns
ns
Alone
In coculture
VIM
CCL2
Relative gene expression to Rhoa
ND
Alone
In coculture
ZEB2

**Supplementary Figure S2. LAD2 and HT-29 cell fractions are 99% and 98% pure after coculture experiments. A**. Scheme of the experimental setup of coculture experiments. **B**. Scatter plot of "LAD2 cell fraction": Cells were labeled with CD45-AF488 and EpCam-AF647 antibodies and analyzed by flow cytometry. **C**. Percentage of CD45-AF488 LAD2 cells and EpCam-AF647 HT-29 cells in "LAD2 cell fraction". N=3. **D**. Scatter plot of "HT-29 cell fraction": cells were labeled and analyzed as in B. **E**. Percentage of CD45-AF488 LAD2 cells and EpCam-AF647 HT-29 cells in "HT-29 cell fraction". N=3. **F**. Relative qRT-PCR of EMT-related markers and CCL2 (left) and ZEB2 (right, not detected (ND)) in HT-29 cells cultured alone or in coculture with BMMCs for 6h. N=3. **G**. Relative qRT-PCR of EMT-related markers and CCL2 (left) and ZEB2 (right, not detected (ND)) in SW403 cells cultured alone or in coculture with LAD2 cells for 6h. N=3. For all panels, lines indicate mean +/- SD and each point represents an independent biological replicate. Significance was determined by two-tailed t-test, $*p \leq 0.05$; $**p \leq 0.01$; $***p \leq 0.001$; $****p \leq 0.0001$, ns- not significant. Supplementary Figure S2A was created with BioRender.com (Lanzloth, R. (2025), https://BioRender.com/k55t245).

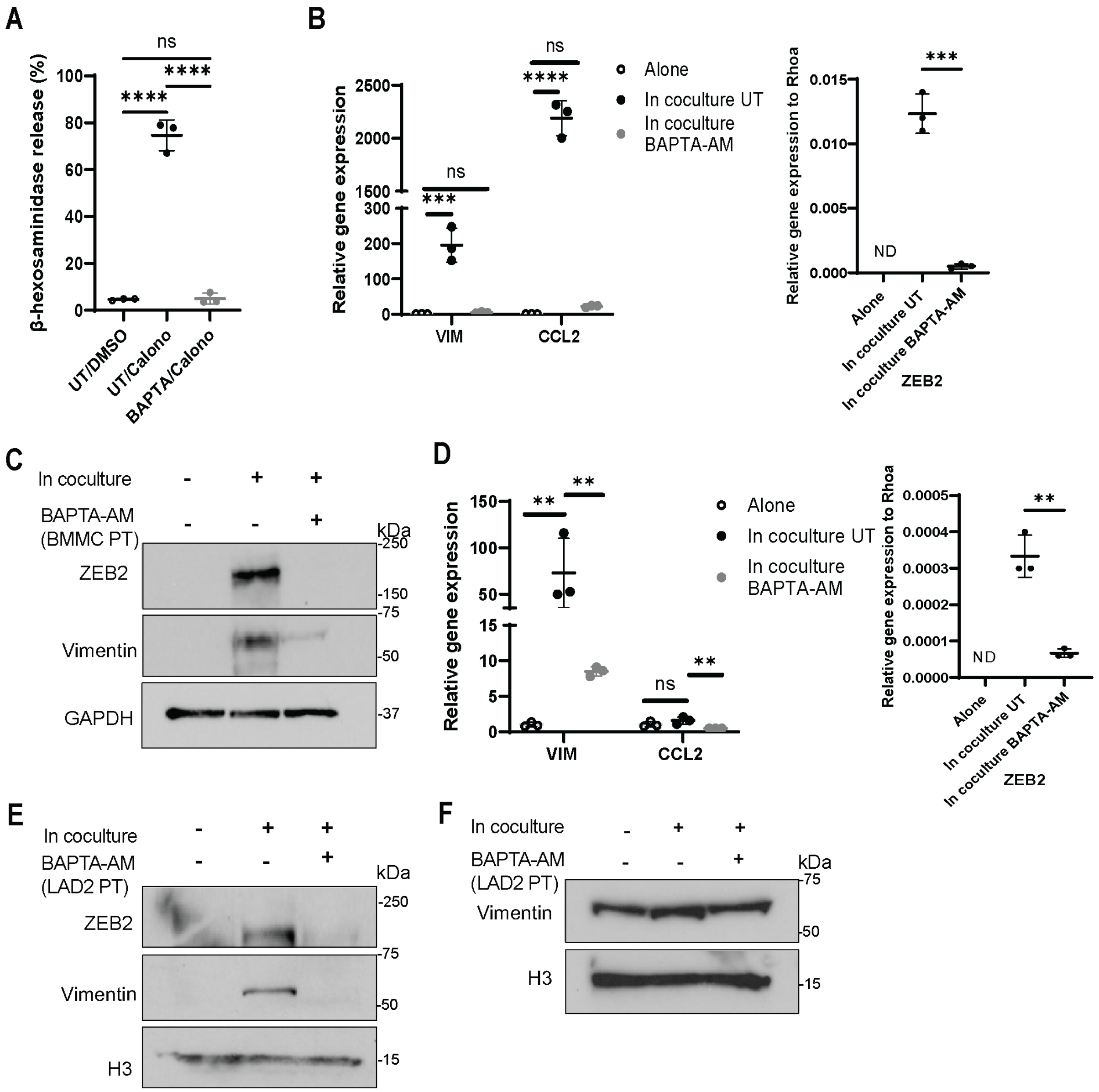
A
ns
****
****
β-hexosaminidase release (%)
UT/DMSO
UT/Calono
BAPTA/Calono
B
Relative gene expression
ns
****
ns
***
VIM
CCL2
Alone
In coculture UT
In coculture BAPTA-AM
Relative gene expression to Rhoa
***
ND
Alone
In coculture UT
In coculture BAPTA-AM
ZEB2
C
In coculture
BAPTA-AM (BMMC PT)
kDa
ZEB2
Vimentin
GAPDH
D
Relative gene expression
**
**
ns
**
VIM
CCL2
Alone
In coculture UT
In coculture BAPTA-AM
Relative gene expression to Rhoa
**
ND
Alone
In coculture UT
In coculture BAPTA-AM
ZEB2
E
In coculture
BAPTA-AM (LAD2 PT)
kDa
ZEB2
Vimentin
H3
F
In coculture
BAPTA-AM (LAD2 PT)
kDa
Vimentin
H3

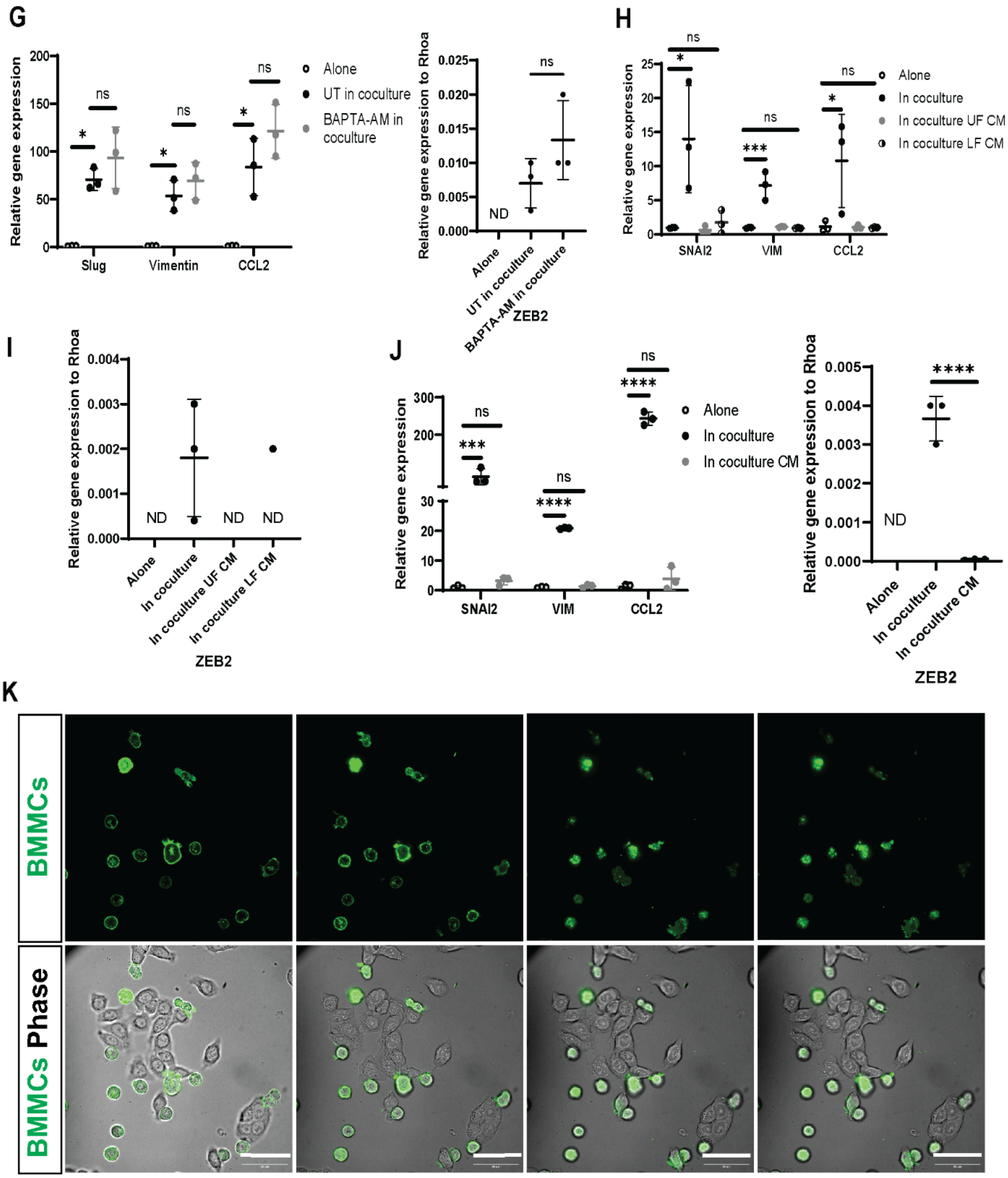


**Supplementary Figure S3. EMT-related marker expression is calcium and contact-dependent in additional CRC cell and MC lines. A**. Percentage of Β-hexosaminidase released in LAD2 cells untreated (UT)/DMSO, untreated in response to calcium ionophore (CaIono; 2 µM, 1h) and pretreated with BAPTA-AM (20 µM, 1h) in response to calcium ionophore. N=1. **B**. Relative qRT-PCR of EMT-related marker genes and CCL2 (left) and

ZEB2 (right, not detected (ND)) in HT-29 cells alone, in coculture with un-pretreated BMMCs, or in coculture with BAPTA-AM (20 μM, for 1h) pretreated BMMCs for 6h. **C**. Western Blot of HT-29 cells alone, in coculture with unpretreated BMMCs, or in coculture with BAPTA-AM pretreated (20 μM, for 1h) BMMCs (BMMC PT) for 3h. N=1. **D**. Relative qRT-PCR of EMT marker genes and CCL2 (left) and ZEB2 (right, not detected (ND)) in SW403 cells alone, in coculture with unpretreated LAD2 cells, or in coculture with BAPTA-AM (20 μM, for 1h) pretreated LAD2 cells for 6h. **E**. Western Blot of SW403 cells alone, in coculture with unpretreated LAD2 cells, or in coculture with BAPTA-AM (20 μM, for 1h) pretreated LAD2 cells for 3h. N=2. **F**. Western blot of LAD2 cells alone, untreated in coculture with HT-29 cells, or BAPTA-AM (20 μM, 1h) pretreated (PT) in coculture with HT-29 cells for 3h. N=3. **G**. Relative qRT-PCR of EMT-related marker genes and CCL2 (left) and ZEB2 (right, not detected (ND)) in HT-29 cells alone, untreated in coculture with LAD2 cells, or pretreated with BAPTA-AM (20 μM, for 1h) in coculture with LAD2 cells for 6h. **H,I**. Relative qRT-PCR of EMT marker genes and CCL2 (H) and ZEB2 (I, not detected (ND)) in HT-29 cells alone, in coculture with LAD2 cells, in concentrated upper fraction of CM (UF CM), or in concentrated lower fraction CM (LF CM) for 6h. **J**. Relative qRT-PCR of EMT marker genes and CCL2 (left) and ZEB2 (right, not detected (ND)) in HT-29 cells alone, in coculture with LAD2 cells, or in coculture conditioned media (CM) for 6h. **K**. BMMCs were labeled with DiO (10 μg/ml, 20 minutes, green) and incubated in direct coculture with HT-29 cells (unlabeled). Each panel represents a different focal plane to better demonstrate cell-cell interactions. Live cells were imaged with Olympus OSR SD confocal microscope for 1h. Magnification 60x. Scale bar = 50 μM. Second picture of panel was also used for Figure 4D. For all panels, lines indicate mean +/- SD and each point represents an independent biological replicate. Significance was determined by two-tailed t-test (B right, D right, F right, G right) and one-way ANOVA (A, B left, D left, F left, G left), *$p \leq 0.05$; **$p \leq 0.01$; ***$p \leq 0.001$; ****$p \leq 0.0001$, ns- not significant.

**Supplementary Video V1. MCs directly interact with CRC cells.** BMMCs were labeled with DiO (10 μg/ml, 20 minutes, green) and incubated in direct coculture with HT-29 cells (unlabeled). Live cells were imaged with Olympus OSR SD confocal microscope. Magnification 60x. Scale bar = 50 μM.

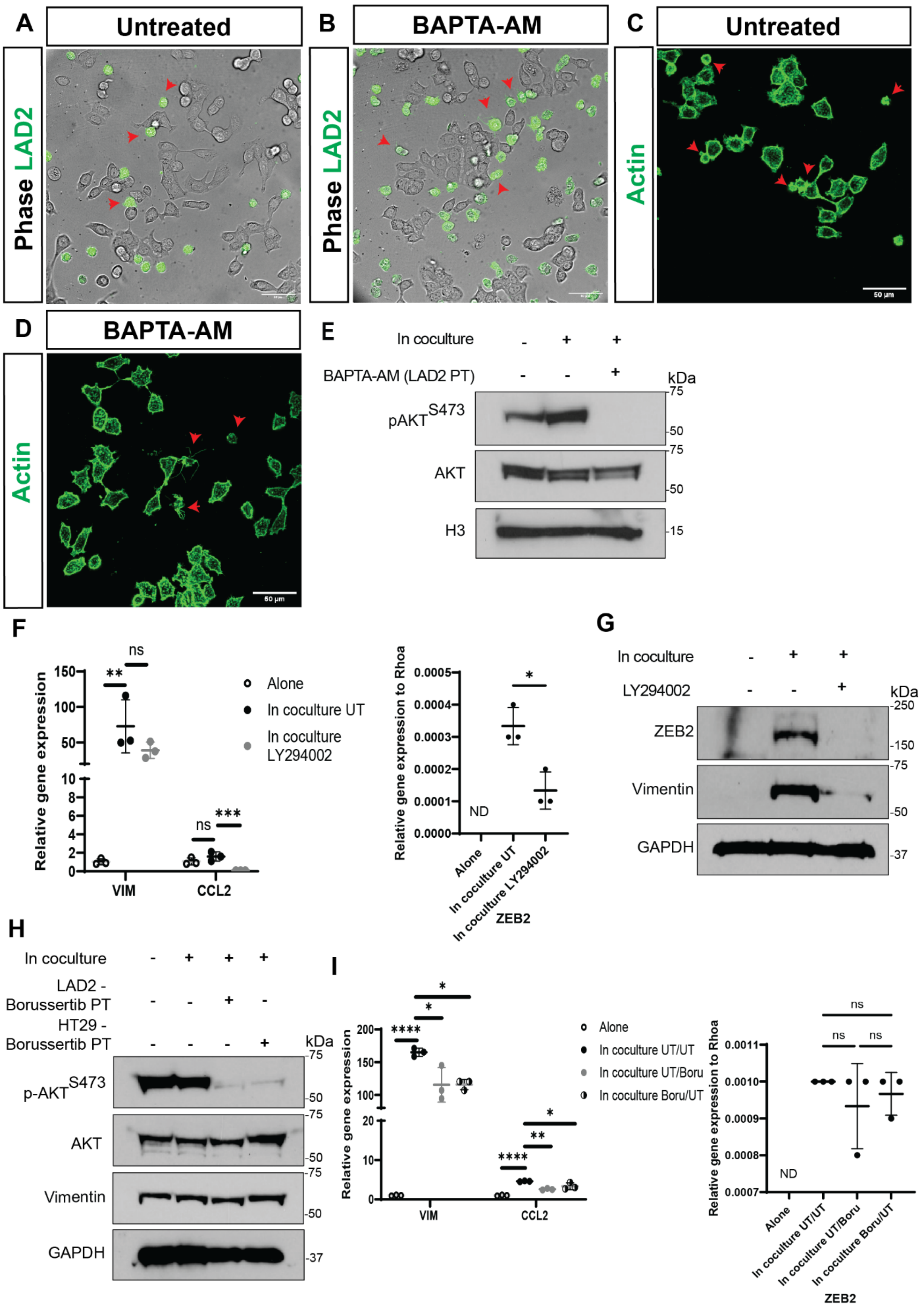

A
Untreated
Phase LAD2
B
BAPTA-AM
Phase LAD2
C
Untreated
Actin
50 µm
D
BAPTA-AM
Actin
50 µm
E
In coculture - + +
BAPTA-AM (LAD2 PT) - - +
kDa
pAKTS473
AKT
H3
75
50
75
50
15
F
Relative gene expression
150
100
50
10
8
6
4
2
0
**
ns
ns
***
VIM
CCL2
Alone
In coculture UT
In coculture LY294002
Relative gene expression to Rhoa
0.0005
0.0004
0.0003
0.0002
0.0001
0.0000
*
ND
Alone
In coculture UT
In coculture LY294002
ZEB2
G
In coculture - + +
LY294002 - - +
kDa
ZEB2
Vimentin
GAPDH
250
150
75
50
37
H
In coculture - + + +
LAD2 - Borussertib PT - - + -
HT29 - Borussertib PT - - - +
kDa
p-AKTS473
AKT
Vimentin
GAPDH
75
50
75
50
75
50
37
I
Relative gene expression
200
150
100
20
10
0
****
*
*
****
**
*
VIM
CCL2
Alone
In coculture UT/UT
In coculture UT/Boru
In coculture Boru/UT
Relative gene expression to Rhoa
0.0011
0.0010
0.0009
0.0008
0.0007
ns
ns
ns
ND
Alone
In coculture UT/UT
In coculture UT/Boru
In coculture Boru/UT
ZEB2

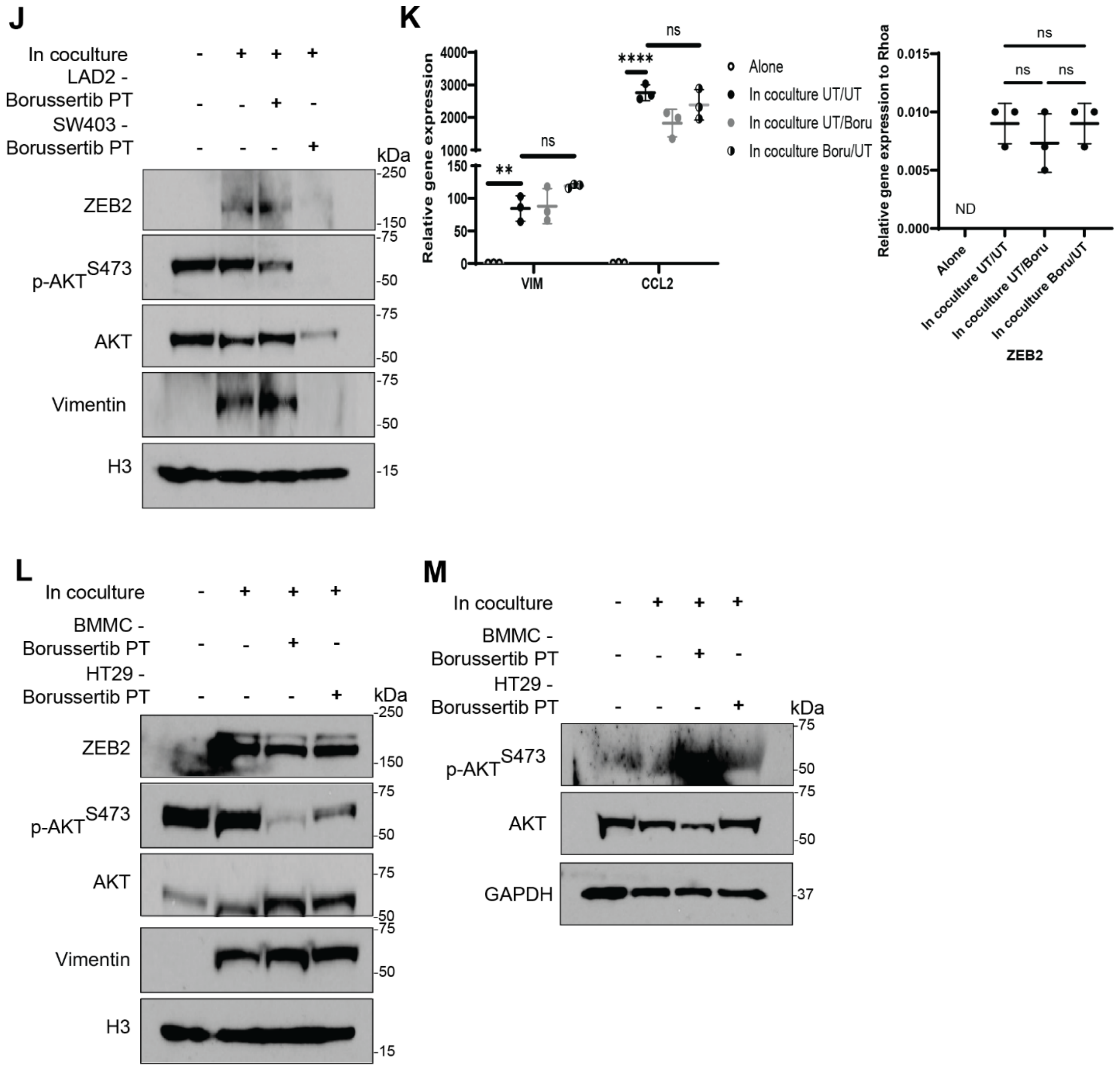


**Supplementary Figure S4. Role of AKT activation in the induction in EMT-related marker expression in CRC cell lines by MCs. A, B**. Untreated LAD2 cells (A) or BAPTA-AM (20 µM, 1h) pretreated LAD2 cells (B) were labeled with DiO (10 µg/ml, 20 minutes, green) and incubated in direct coculture with HT-29 cells (unlabeled). Live cells were imaged with Olympus OSR SD confocal microscope for 3h. Magnification 40x. **C,D**. Untreated LAD2 cells (C) or BAPTA-AM (20 µM, 1h) pretreated LAD2 cells (D) were incubated in direct coculture with HT-29 cells. Actin cytoskeleton was labeled with Phalloidin-488 reagent. Live cells were imaged with Leica Stellaris 8 FALCON confocal microscope. Magnification 63x. **E**. Western blot of LAD2 cells alone, untreated in coculture with HT-29 cells, or BAPTA-AM (20 µM, 1h) pretreated (PT) in coculture with HT-29 cells for 3h. Western blots for Supplementary Figure S4E and S3H originate from the same blot and have the same loading control. N=3. **F**. Relative qRT-PCR of EMT related genes and CCL2 (left) and ZEB2 (right, not detected (ND)) in SW403 cells alone, in coculture with untreated LAD2 cells, or in coculture with LAD2 cells under LY294002 treatment (60 μM) for 6h. **G**. Western blot of SW403 cells treated and in cocultured as in F for 3h. N=1. **H**. Western blot of LAD2 cells unpretreated and alone, unpretreated and in coculture with unpretreated HT-29 cells, Borussertib (1 μM, overnight) pretreated in coculture with unpretreated HT-29 cells (LAD2 – Borussertib PT), unpretreated in coculture with Borussertib (10 μM, overnight) pretreated HT-29 cells (HT-29 – Borussertib PT) for 3h. N=3. **I**. Relative qRT-PCR of EMT related genes and CCL2 (left) and ZEB2 (right, not detected (ND)) in SW403 cells alone, unpretreated (UT) in coculture with unpretreated LAD2 cells (UT/UT), or unpretreated in coculture with Borussertib (1 μM, overnight) pretreated LAD2 cells (UT/Boru), or Borussertib (10 μM, 1h) pretreated in coculture with unpretreated LAD2 cells (Boru/UT) for 6h. **J**. Western blot of SW403 cells treated and cocultured as in I. N=2. **K**. Relative qRTP-PCR of EMT related genes and CCL2 (left) and ZEB2 (right, not detected (ND)) in HT-29 cells unpretreated and alone, unpretreated and in coculture with unpretreated BMMCs (UT/UT), unpretreated in coculture with Borussertib (1 μM, overnight) pretreated BMMCs (UT/Boru), Borussertib (10 μM, overnight) pretreated in coculture with unpretreated BMMCs (Boru/UT) for 6h. **L**. Western blot of HT-29 cells treated and cocultured as in K for 3h. N=1. **M**. Western blot of BMMCs treated and cocultured as in H. N=1. For all panels, lines indicate mean +/- SD and each point represents an independent biological replicate. Significance was determined by one-way ANOVA (F , I, K), *$p \leq 0.05$; **$p \leq 0.01$; ***$p \leq 0.001$; ****$p \leq 0.0001$, ns- not significant.

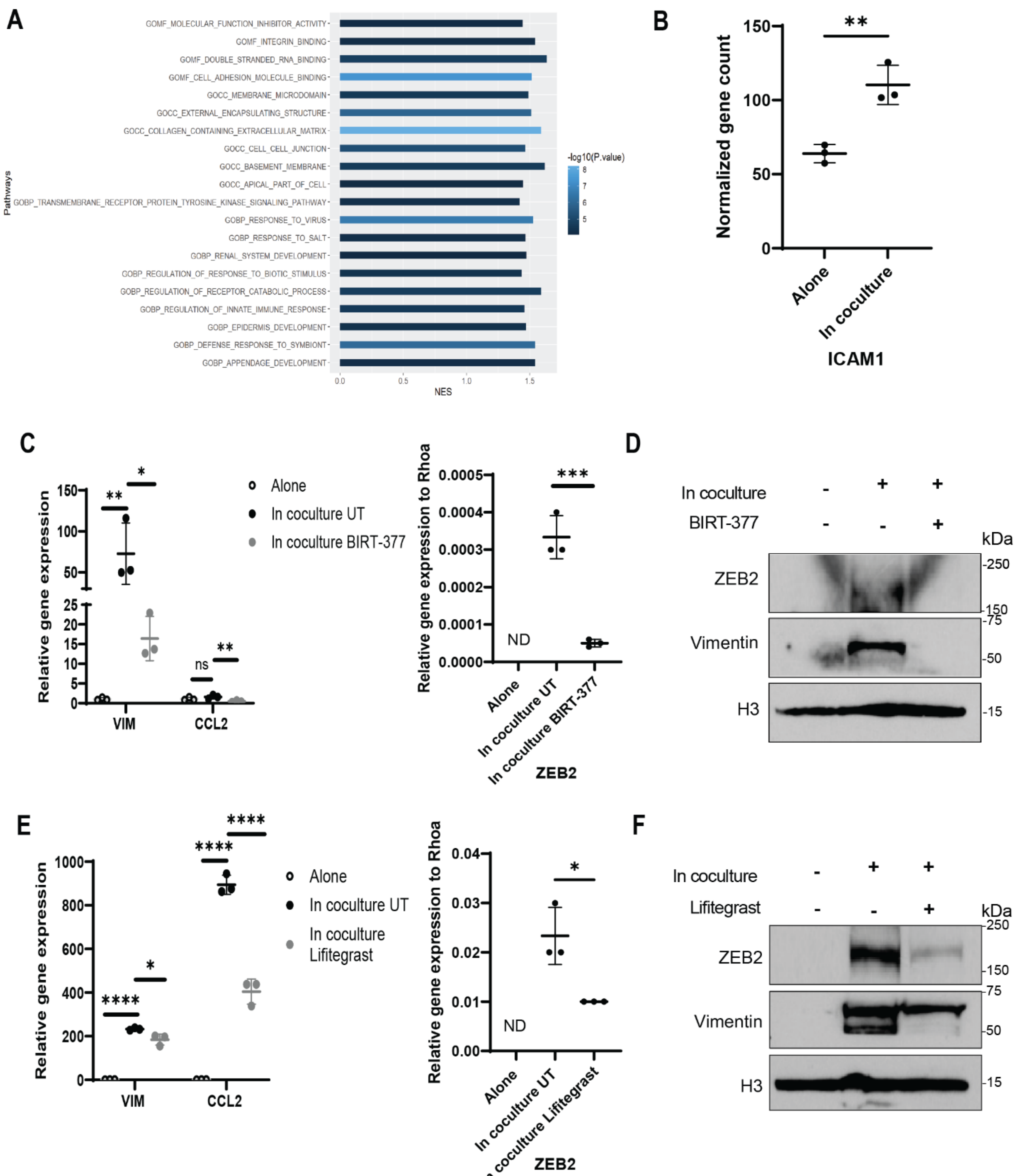


**Supplementary Figure S5. LFA-1/ICAM-1 integrins are involved in the induction of the EMT-related marker expression in additional CRC cells. A**. Gene ontology enrichment of RNA-sequencing data from LAD2 cells cocultured with HT-29 cells for 12h. **B**. Normalized gene count of *ICAM1* gene from RNA-sequencing data from empty vector HT-29 cells alone or in coculture with LAD2 cells for 12h. **C**. Relative qRT-PCR of EMT related genes and CCL2 (left) and ZEB2 (right, not detected (ND)) in SW403 cells alone, in coculture with untreated

(UT) LAD2 cells, or in coculture with LAD2 cells under BIRT-377 treatment (in coculture BIRT-377) (40 μM) for 6h. **D**. Western blot of SW403 cells treated and cocultured as in C for 3h. N=1. **E**. Relative qRT-PCR of EMT related genes and CCL2 (left) and ZEB2 (right, not detected (ND)) in HT-29 cells alone, in coculture with untreated (UT) BMMCs, or in coculture with BMMCs under Lifitegrast treatment (40 μM) for 6h. **F**. Western blot of HT-29 cells treated and cocultured as in E for 3h. N=1. For all panels, lines indicate mean +/- SD and each point represents an independent biological replicate. Significance was determined by one-way ANOVA (C and E), *$p \leq 0.05$; **$p \leq 0.01$; ***$p \leq 0.001$; ****$p \leq 0.0001$, ns- not significant.

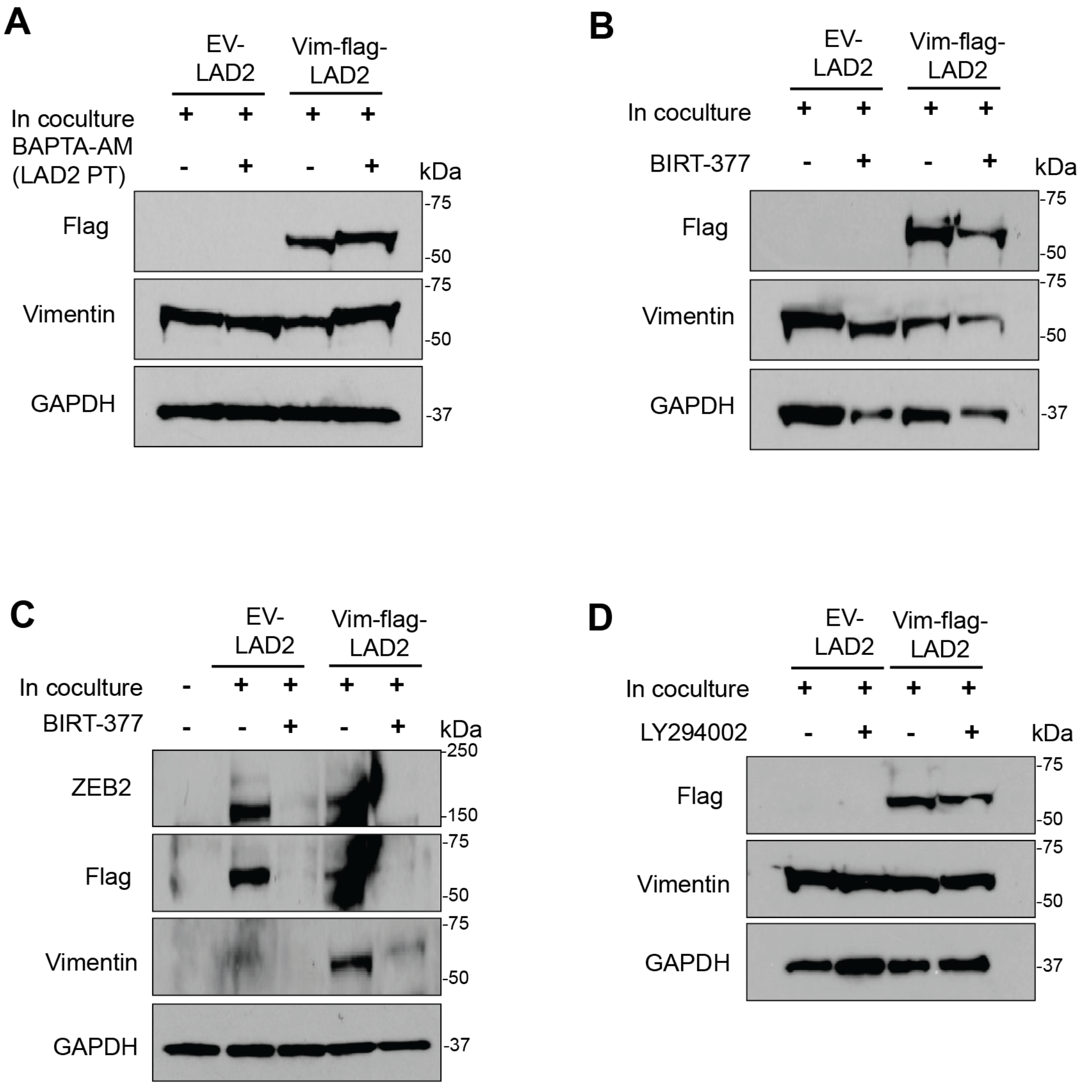


**Supplementary Figure S6. Treatment does not change Vimentin-Flag expression in MCs. A**. Western blot of unpretreated empty vector (EV-) LAD2 cells in coculture with HT-29 cells, BAPTA-AM (20 μM, 1h) pretreated EV LAD2 cells in coculture with HT-29 cells, unpretreated Vimentin-Flag transduced (Vim-Flag-) LAD2 cells in coculture with HT-29 cells, or BAPTA-AM (20 μM, 1h) pretreated Vim-Flag LAD2 cells in coculture with HT-29 cells for 3h. N=1. **B**. Western blot of untreated EV LAD2 cells in coculture with HT-29 cells, EV LAD2 cells in coculture with HT-29 cells under BIRT-377 treatment (20 μM), untreated Vim-Flag LAD2 cells in coculture with HT-29 cells, or Vim-Flag LAD2 cells in coculture with HT-29 cells under BIRT-377 treatment (20 μM) for 3h. N=1. **C**. Western blot of SW403 cells alone, in coculture with untreated EV LAD2 cells, in coculture with EV LAD2 cells under BIRT-377 treatment (40μM), in coculture with untreated Vim-Flag LAD2 cells, or in coculture with Vim-Flag LAD2 cells under BIRT-377 treatment (40μM) for 3h. N=1. **D**. Western blot of untreated EV LAD2 cells in coculture with HT-29 cells, EV LAD2 cells in coculture with HT-29 cells under LY294002 treatment (50 μM), untreated Vim-Flag LAD2 cells in coculture with HT-29 cells, or Vim-Flag LAD2 cells in coculture with HT-29 cells under LY294002 treatment (50 μM) for 3h. N=1.
